# A chromosome-level genome assembly of the Hispid cotton rat (*Sigmodon hispidus*), a model for human pathogenic virus infections

**DOI:** 10.1101/2024.03.21.586163

**Authors:** Jingtao Lilue, André Corvelo, Jèssica Gómez-Garrido, Fengtang Yang, Keiko Akagi, Gia Green, Bee Ling Ng, Beiyuan Fu, Uciel Pablo Chorostecki, Sarah Warner, Marina Marcet-Houben, Thomas Keane, James C. Mullikin, Tyler Alioto, Toni Gabaldón, Benjamin Hubert, David E. Symer, Stefan Niewiesk

## Abstract

**Background:** The cotton rat (*Sigmodon hispidus*), a rodent species native to the Americas, has emerged as a valuable laboratory model of infections by numerous human pathogens including poliovirus and respiratory syncytial virus (RSV).

**Results:** Here we report the first reference assembly of the cotton rat genome organized at a chromosomal level, providing annotation of 24,878 protein-coding genes. Data from PCR-free whole genome sequencing, linked-read sequencing and RNA sequencing from pooled cotton rat tissues were analyzed to assemble and annotate this novel genome sequence. Spectral karyotyping data using fluorescent probes derived from mouse chromosomes facilitated the assignment of cotton rat orthologs to syntenic chromosomes, comprising 25 autosomes and a sex chromosome in the haploid genome. Comparative phylome analysis revealed both gains and losses of numerous genes including immune defense genes against pathogens. We identified thousands of recently retrotransposed L1 and SINE B2 elements, revealing widespread genetic innovations unique to this species.

**Conclusions:** We anticipate that annotation and characterization of the first chromosome-level cotton rat genome assembly as described here will enable and accelerate ongoing investigations into its host defenses against viral and other pathogens, genome biology and mammalian evolution.

## INTRODUCTION

Cotton rats (*Sigmodon hispidus*) are small rodents extant in the New World. They live in the southern part of the United States, Central America and the northern part of South America (Prince 1994)(Niewiesk and Prince 2002)(Green, Huey and Niewiesk 2013). These rodents have been used as laboratory animals since the 1930’s, when in pursuit of a non-primate animal model they were infected with poliovirus (Armstrong 1939). The cotton rat subsequently has been found to be susceptible to a large number of human pathogens, and has been used in studies of the pathogenesis of human respiratory viral infections and in testing of vaccines and antivirals (Niewiesk and Prince 2002)(Green, Huey and Niewiesk 2013). Many human pathogens replicate well in cotton rat but not in house mouse (*Mus musculus*), including RSV, human metapneumovirus, influenza virus and adenovirus. RSV replicates approximately one hundred-fold more in cotton rats, and as a consequence, model studies to test the efficacy of vaccines in humans provide a better predictive value in cotton rat than in mouse (Prince et al. 1978)(Boukhvalova, Prince and Blanco 2009)(Cullen, Blanco and Morrison 2015). The cotton rat is a natural carrier of additional respiratory viruses with tropism for humans, including the Black Creek Canal virus strain of hantavirus. Moreover, other viral pathogens infecting the respiratory tract in humans do not replicate at all in mouse, including parainfluenza virus and measles virus (Green, Huey and Niewiesk 2013).

A comprehensive approach to the analysis of overall immune responses to infection and vaccination in cotton rats requires measurements of gene expression profiles. However, the lack of a cotton rat reference genome has limited such experiments. In recent studies, immune responses of RSV-infected cotton rats were evaluated by comparing RNA expression patterns in infected versus uninfected lungs (Rajagopala et al. 2018) (Strickland et al. 2022). As in humans, numerous immune genes related to defense against viral infections were upregulated, while others were downregulated late during infection. Although these results indicate broad similarities with gene expression changes in humans as quantified by microarray (Mejias et al. 2013)(Heinonen et al. 2020), additional cotton rat genes appeared novel, as no known orthologs have been identified in other species.

We anticipated that development of an annotated cotton rat reference genome would facilitate future investigations into the complex interplay between infectious pathogens and host immune responses. In addition, a high quality, annotated reference assembly would enable comprehensive comparisons of immune responses across species such as between cotton rat and human, for example through the development of agnostic assays of cotton rat gene expression including RNA-seq, single cell RNA-seq and proteomics. To develop such an annotated cotton rat genome assembly, we sequenced cotton rat DNA and pooled RNA isolated from several tissues. Our analysis defined the cotton rat genome length and chromosomes, annotated genes including immune genes and their orthologs, identified active, novel retrotransposon families, and assessed phylogeny. The resulting draft reference genome assembly of the cotton rat has highlighted a few dozen candidate genes for their potential roles in host defenses against various viruses. We anticipate that this new cotton rat genome assembly will facilitate future research on pathogenic infections, host immunity, genome biology, mammalian evolution and potentially additional research fields.

## RESULTS

**Genome assembly.** Genomic DNA extracted from a male cotton rat was sequenced at 72x depth of coverage from linked reads (10x Genomics Chromium), 38x depth of coverage with a PCR-free whole genome sequencing (WGS) library with ∼350 nucleotide (nt) mean genomic DNA insert length (Illumina), and 40x depth of coverage with a second PCR-free WGS library with ∼550 nt mean insert length.

To assemble a high-quality draft of the cotton rat genome, we combined the linked-read sequencing data and WGS data from the two PCR-free libraries. First, error-corrected linked reads were assembled using Supernova v2.0.1 (Weisenfeld et al. 2017) to produce a draft pseudo-haploid representation of the genome. We used ARKS v1.0.2 (Coombe et al. 2018) to scaffold the draft further, resulting in a ∼2.50 Gb assembly with a scaffold N50 length of 4.95 megabasepairs (Mb). Assembly gaps were closed by using error-corrected PCR-free short-read data in two complementary strategies (see Materials and Methods), leading to a significant increase in contig N50 lengths which improved from the original 62.8 kilobasepairs (kb) to 360.5 kb (**Table 1**).

**Table 1.** *S. hispidus* genome assembly statistics.

| metric | Supernova<br>v2.01 | Supernova<br>v2.0.1 + ARKS | Supernova v2.0.1 +<br>ARKS + gap closure<br>+ redundancy<br>removal | SigHis_v1_BIUU<br>GenBank:<br>GCA_004025045.1BioSample:<br>SAMN07678138 |
| --- | --- | --- | --- | --- |
| length (bp) | 2,514,336,970 | 2,514,367,750 | 2,504,610,684 | 2,730,600,022 |
| scaffolds (N) | 23,680 | 20,602 | 17,569 | 883,546 |
| longest<br>scaffold (bp) | 7,913,074 | 24,396,235 | 24,371,482 | 1,032,205 |
| scaffold N50<br>length (bp) | 1,572,529 | 4,921,260 | 4,950,377 | 101,373 |
| scaffold N90<br>length (bp) | 209,288 | 746,302 | 784,357 | 1,886 |
| contigs (N) | 87,086 | 87,086 | 31,279 | 897,290 |
| longest contig<br>(bp) | 520,409 | 520,409 | 2,485,181 | 783,669 |
| contig N50<br>length (bp) | 62,801 | 62,801 | 360,531 | 67,983 |
| contig N90<br>length (bp) | 16,655 | 16,655 | 78,531 | 1,879 |
| gaps (N) | 63,415 | 66,492 | 13,710 | 13,744 |
| gapped length<br>(bp) | 29,394,360 | 29,425,140 | 23,026,440 | 1,374,400 |

The resulting assembly shows a high level of sequence completeness, with 99.28% of PCR-free library reads mapping to it, and 98.69% aligned in proper pairs. This high level of completeness is also corroborated by a high degree of observed gene-space completeness, with 242 out of 248 (97.6%) core eukaryotic genes and 8,919 out of 9,226 (96.7%) mammalian ortholog groups identified in their complete forms, as determined by CEGMA (Parra, Bradnam and Korf 2007) and BUSCO (Simao et al. 2015), respectively.

Another independent genome sequence dataset for *S. hispidus* recently was deposited at the National Center for Biotechnology Information (NCBI), under assembly name SigHis_v1_BIUU (GenBank accession number GCA_004025045.1; BioSample SAMN07678138; BioProject PRJNA399433). To compare various features of this recent dataset with ours, we tabulated several of its reported parameters, including total sequenced genome length, number of scaffolds, scaffold N50 length, and number of gaps (**Table 1**). Its depth of sequencing coverage genomewide was 43.2x, substantially less than the cumulative coverage of 150x reached in our study. The SigHis_v1_BIUU assembly has an approximately 50-fold higher count of single-contig scaffolds, and its scaffold N50 lengths are approximately 40-fold lower (**Table 1**). Its cumulative gap length is less than that in our genome assembly, but this is likely attributable to an arbitrary assumption of only 100 nt per gap as reported in that assembly. To our knowledge, chromosomes have not been assigned to date in the BIUU assembly.

### Cross-species chromosome painting and assembly

To assign draft chromosome models for the *S. hispidus* genome sequence assembly, we performed comparative chromosome painting of metaphase spreads, using probes derived from flow-sorted autosomes and the X chromosome of the mouse, *Mus musculus* (**Supp. Fig. S1**). *S. hispidus* chromosomes were identified and numbered based on G-banded metaphases (Elder 1980). As expected, *S. hispidus* shows extensive synteny when compared with the mouse chromosomes over long genomic distances (**Supp. Fig. S2**). Using the mouse-derived chromosome paints, a strong, continuous signal was observed across most portions of *S. hispidus* chromosomes. However, some small regions of the chromosomes were ambiguous, either displaying multiple probes derived from multiple mouse chromosomes (e.g. genomic repeats) or no probe signal. This result confirmed the accumulation of considerable chromosome-level differences between the two rodent species, despite very high overall levels of orthology and synteny that were observed.

We assembled the *de novo* draft cotton rat genome sequence scaffolds into 26 pseudo-chromosomes, based on chromosome paints data and on their synteny and orthologous sequence similarities with mouse. The set of pseudo-chromosome sequences contains 803 scaffolds, with a total length of 2.359M basepairs covering approximately 92.1% of the draft haploid *de novo* cotton rat assembly. These data indicate that the karyotype of *S. hispidus* is similar to an ancestral karyotype of the genus *Sigmodon*, which harbors 2n = 52 chromosomes (i.e. counting both autosomes and sex chromosomes). This finding also corroborates a previous report indicating that the cotton rat genome has accumulated few or no changes in chromosome counts or composition when compared with five other *Sigmodon* species (*S. hirsutus, S. leucotis, S. ochrognathus, S. peruanus*, and *S. toltecus*) (Swier et al. 2009).

These findings of orthologous sequences and synteny among distinct rodent species were corroborated further upon comparisons with previously reported chromosome assignments in a New World mouse species, *Onychomys torridus* (*O. torridus*, also known as the Southern grasshopper mouse), and in the white-footed mouse (*Peromyscus leucopus*) (Long et al. 2019), based on similar methods (**Supp. Fig. S3, Supp. Fig. S4, Figure 1**). The results demonstrate a high degree of interspecific relatedness when comparing cotton rat vs. these independent rodent species.

**Figure 1.**
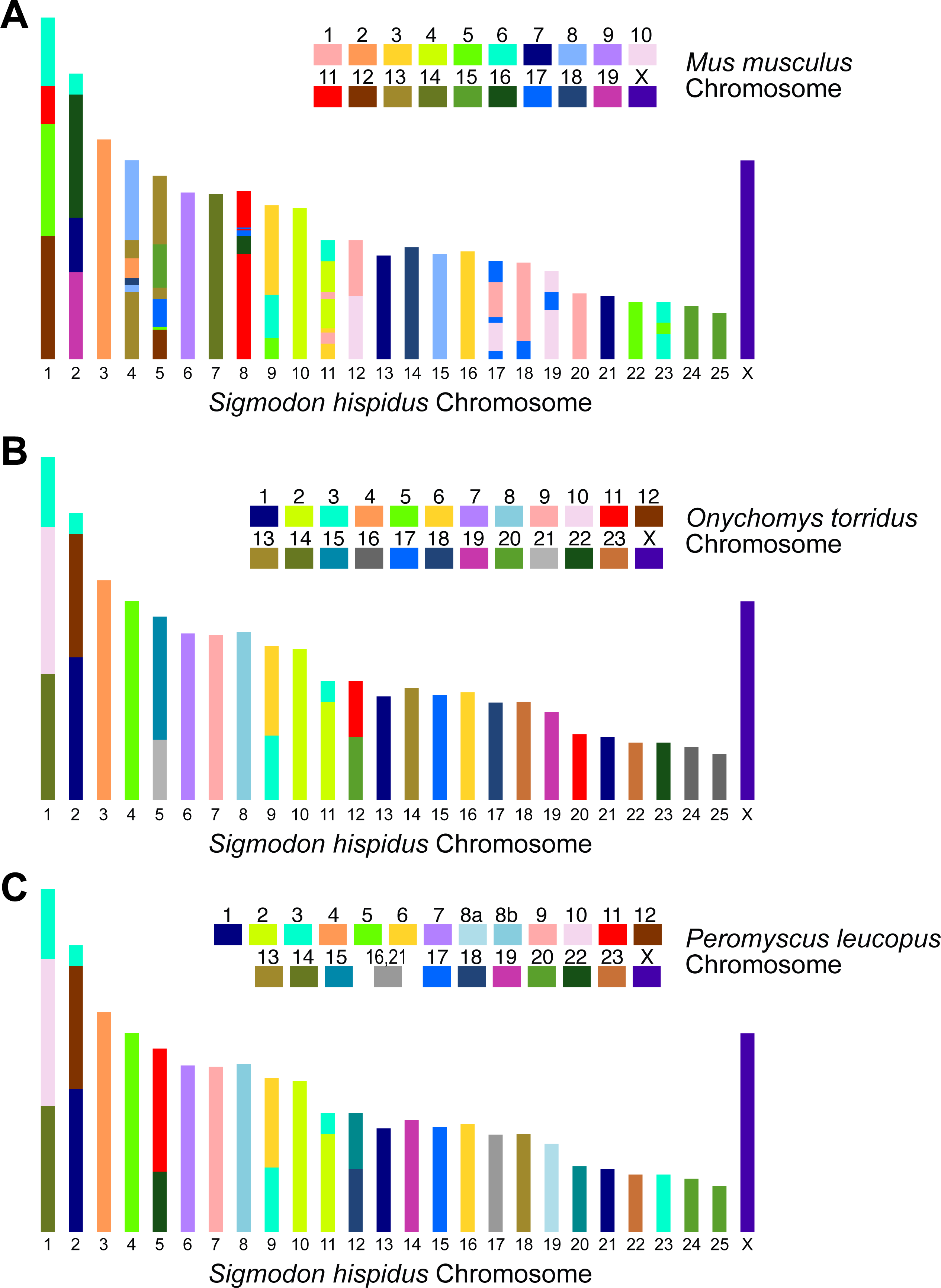
Comparison of pseudo-chromosome assemblies of cotton rat (*S. hispidus*), determined experimentally with mouse chromosome paints or upon alignments with *O. torridus* and *P. leucopus* orthologous segments. Schematics represent karyotypes from metaphase spreads of cotton rat (*S. hispidus*), (A) painted with probes derived from *M. musculus* chromosomes (*key*) or annotated from alignments with syntenic segments from annotated pseudo-chromosomes of (B) *O. torridus* (probed with mouse chromosome paints, cf. Methods) and (C) *P. leucopus* (Long et al. 2019). Color codes on *S. hispidus* chromosomes are based on *Mus musculus*.

### Cotton rat genes

Combining outputs from three *ab initio* gene-calling programs (see **Methods**), we identified 24,878 protein-coding genes across the cotton rat genome assembly. From these genes, we conservatively predicted expression of a total of 29,010 transcripts (i.e. 1.17 transcripts per gene), while ignoring most other potential isoforms arising from alternative splicing. In turn, these transcripts are predicted to encode 28,403 unique protein products. We assigned functional labels to 90.8% of the predicted gene products. On average, each annotated gene contains 9.46 exons, with 75% of transcripts predicted as multi-exonic (**Table 2**). In addition, we predicted 40,481 non-coding transcripts, of which 23,669 and 16,812 appear to be expressed from long and short non-coding RNA genes, respectively.

**Table 2.** Summary of annotated protein-coding genes in cotton rat genome assembly.

|  | **Hispid2A annotation** |
| --- | --- |
| number of protein-coding genes | 24,878 |
| median gene length in genome (bp) | 8,808 |
| number of transcripts | 29,005 |
| number of exons | 235,349 |
| number of coding exons | 227,132 |
| exons/transcript | 9.93 |
| transcripts/gene | 1.17 |
| multi-exonic transcripts | 0.75 |
| gene density (gene/Mb) | 9.93 |

We sought to obtain experimental evidence confirming that a large portion of the predicted cotton rat transcriptome is expressed. In addition, concordance between expressed transcript structures and our *ab initio* gene predictions needed to be checked. For these reasons, we performed RNA-seq on two pools of total RNAs. RNA was extracted from multiple tissues extracted from two adult male cotton rats, it was pooled to represent roughly similar concentrations across the tissues from each individual, and finally two RNA-seq libraries were prepared and sequenced. One of the cotton rat individuals was untreated and healthy (naive), while the other was injected intraperitoneally with house dust mite (HDM) antigen, followed by intranasal exposure after eight days and subsequent euthanasia four days thereafter. RNA-seq data from each individual were aligned using STAR v-2.5.3a (Dobin et al. 2013) against our reference assembly genome (**Table 3; Supp. Table S1**).

**Table 3.** Features of RNA-seq read alignments. RNA-seq was conducted on RNA pools from several tissues extracted from naive vs. house dust mite (HDM)-treated cotton rat individuals. Results from these two tissue pools were combined.

| **Read counts and features** | **treatment** |  |  |
| --- | --- | --- | --- |
|  | **naïve** | **HDM** | **combined** |
| RNA-seq read counts | 196,659,156 | 218,393,360 | 415,052,516 |
| % uniquely mapped | 81.69% | 85.84% | 83.77% |
| % mapped to multiple loci | 5.25% | 7.00% | 6.13% |
| % unmapped | 12.89% | 6.87% | 9.88% |
| expressed genes identified | 20,834 | 20,506 | 21,417 |
| % of all annotated protein-coding genes | 93% | 92% | 96% |

At least one RNA-seq transcript read from either pool could be assigned to each of 21,417 genes, which comprise 86.1% of the predicted protein-coding genes (**Table 3; Supp. Fig. S5**). These included 911 genes expressed in the naive individual tissue pool, and 583 in the other pool from the individual exposed to house dust mite (HDM) (**Supp. Fig. S5**). To assess if annotations of our reference genome included immune response genes that were expressed in these pools, first, we selected 224 genes identified in our reference assembly (Supp. Table S2) that were annotated by the specific gene ontology (GO) term “immune response” (GO:0006955). Of these, 213 (95%) were found to be expressed in our RNA-seq data (including 208 expressed genes in the naïve dataset and 207 in the HDM one, out of the total counts in **Table 3**). The 11 remaining “immune response” genes lacking transcripts detected in our RNA-seq pools were tabulated as well (**Supp. Table S3**).

A recent report identified host response transcripts that were differentially expressed upon infection with RSV (Rajagopala et al. 2018). We downloaded a list of their cDNA sequences and aligned them against our reference genome assembly. Of 19 differentially expressed genes that also were annotated and confirmed in our reference assembly, we detected expression for 15 (79%) of them in at least one of our two RNA-seq pools (**Supp. Table S4**).

To investigate structure and expression of key immune gene family members involved in cotton rat responses against RSV and other infectious pathogens, we aligned RNA-seq reads against our reference genome assembly. We focused analysis on clusters of genes in the *IFIT*, *CXCL* and *GBP* families, which have been shown to play important roles in RSV replication. These gene clusters are located on cotton rat Chrs. 2, Chr. 1 and Chr. 16, respectively. No gaps were identified in any of the genes in these gene clusters, highlighting the high quality of our reference genome assembly (**Fig. 2**; **Table 1**). By contrast, when the same RNA-seq reads were aligned against the previously reported genome assembly SigHis_v1_BIUU, numerous gaps interrupting the clustered family members were observed (data not shown).

**Figure 2.**
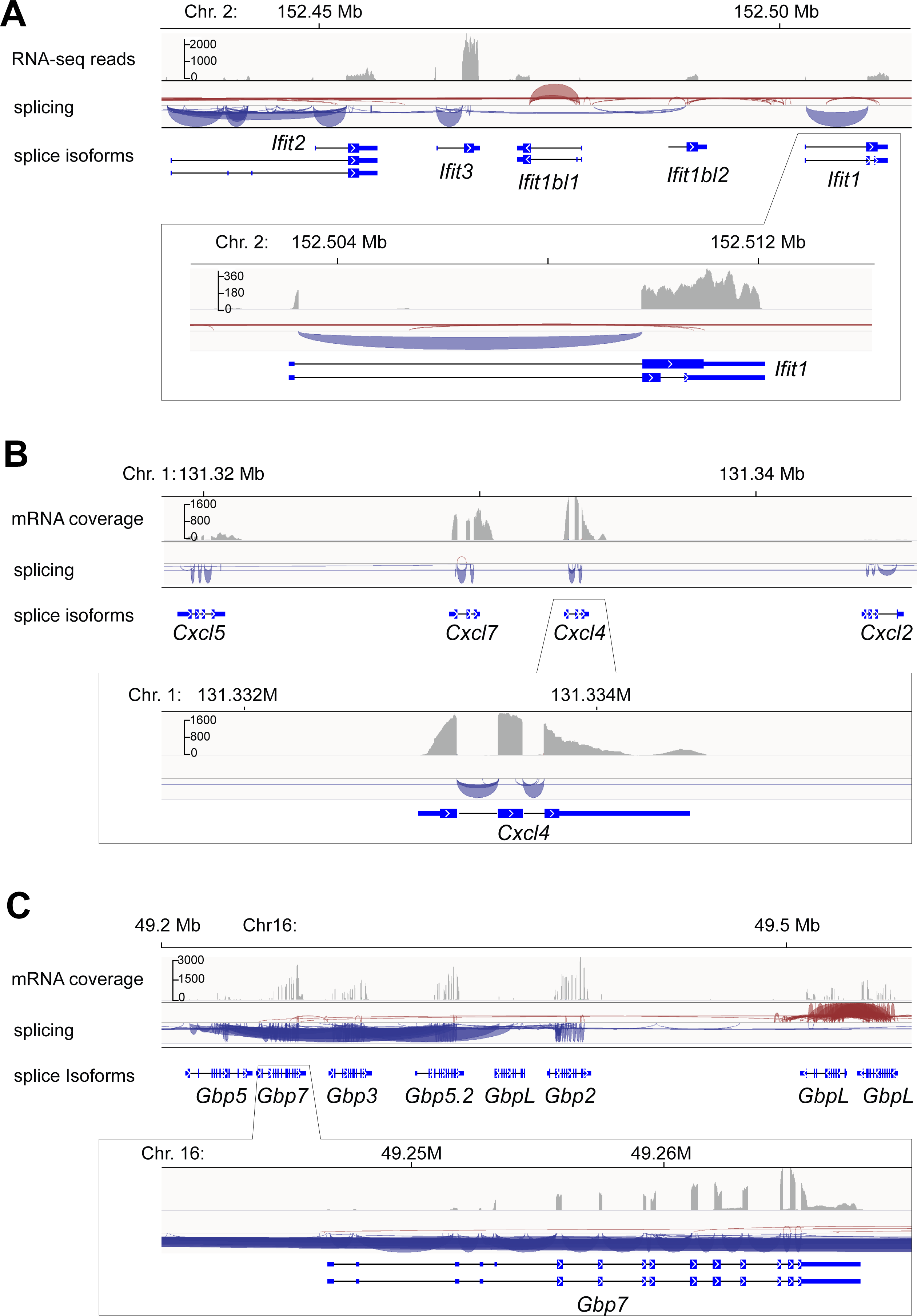
RNA-seq of pooled cotton rat tissues reveals expression patterns of gene family clusters involved in key immune response pathways. Poly(A)-positive mRNAs were extracted from multiple tissues of a healthy donor cotton rat and of a second individual that was exposed to house dust mite antigen, pooled, and reverse transcribed for preparation of strand-specific RNA-seq libraries. Approximately 50 million sequencing read pairs (2×150 bp paired end) were obtained per pool. Sequences were aligned to the refined reference cotton rat genome and compared with gene annotation models. Shown in tracks are: *top*, counts of aligned reads; *middle*, Sashimi plots showing RNA splicing; and *bottom*, gene model schematics depicting exons and introns, at several immune gene family loci including the (A) *Ifit1* gene family (Chr. 2); (B) *Cxcl* gene family (Chr. 1); and (C) *Gbp7* gene (Chr. 16). We detected transcript isoform expression at each annotated gene and multiple instances of alternative splicing. For comparison, recently reported, independent RNA-seq reads from cotton rat individuals exposed to respiratory syncytial virus infection also were aligned to our annotated cotton rat genome assembly (**Supp. Fig. S5**), revealing comparable splicing isoforms but at lower sequencing coverage.

Predicted exons, introns and splice sites were confirmed by the RNA-seq read alignments. We detected expression of alternative splicing isoforms of *Ifit2*, *Ifit1bl1*, *Ifit1* and *Gbp7*, while no splicing isoforms were identified for other family members in the clusters. Counts of RNA-seq reads aligned against each annotated family member in the three families’ chromosomal loci were wide-ranging, but each gene (and predicted exon) was expressed at a detectable level (transcripts per million, TPM > 1).

### Phylome analysis

To investigate the molecular evolution of the cotton rat genome in the context of other sequenced rodents and additional, more distant mammalian relatives, we reconstructed its phylome, i.e. a complete collection of gene evolutionary histories (Fuentes et al. 2022). To make detailed comparisons, we also reconstructed both the mouse and human phylomes based on the same overall set of species. A total of 64,415 maximum likelihood gene phylogenies were reconstructed from the phylomes of these three focus species. To provide an evolutionary framework for our comparisons, we reconstructed the evolutionary relationships between the considered species by concatenating the alignments of 895 widespread genes with one-to-one orthologs in all species, and reconstructing a maximum likelihood tree using RAxML (Stamatakis 2014). The species tree shows the expected topology (**Fig. 3**).

**Figure 3.**
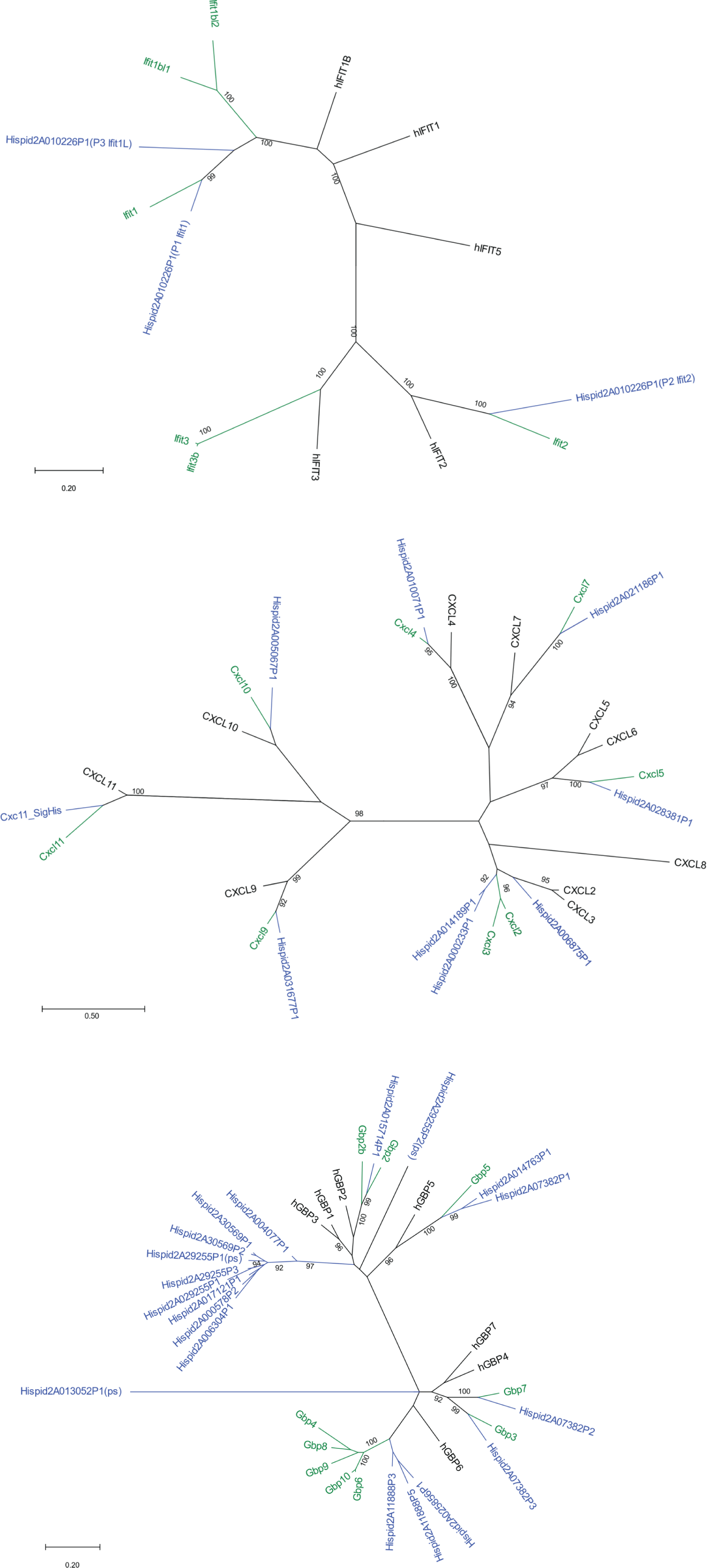
Phylome analysis of key immune response gene families in cotton rat vs. mouse and human genomes. Analysis of gene relatedness was conducted to construct phylome trees, revealing relatedness of individual gene family members from the (A) *Ifit*; (B) *Cxcl* and (C) *Gbp7* families detected in the (blue font) cotton rat, (green) mouse and (black) human genomes. Gene annotation IDs and models for cotton rat genes are available at https://denovo.cnag.cat/cottonrat. *Numbers*, percentage of support for each node.

Next, we examined gene trees in the phylomes to detect gene duplication events (Huerta-Cepas and Gabaldon 2011) that occurred in one or more of the three focus species (i.e. cotton rat, mouse, human). These duplication frequencies are not normalized for time, as we did not develop or use a dated tree. Both rodent species harbor an approximately two-fold increase in the number of species-specific duplications when compared to human, with an average of 0.37 duplications per gene in cotton rat and 0.31 per gene in mouse, compared with just 0.16 in human. This difference may be explained by different divergence times for these focus species from their most recent common ancestor, as represented by branches in the species tree (**Fig. 4**).

**Figure 4.**
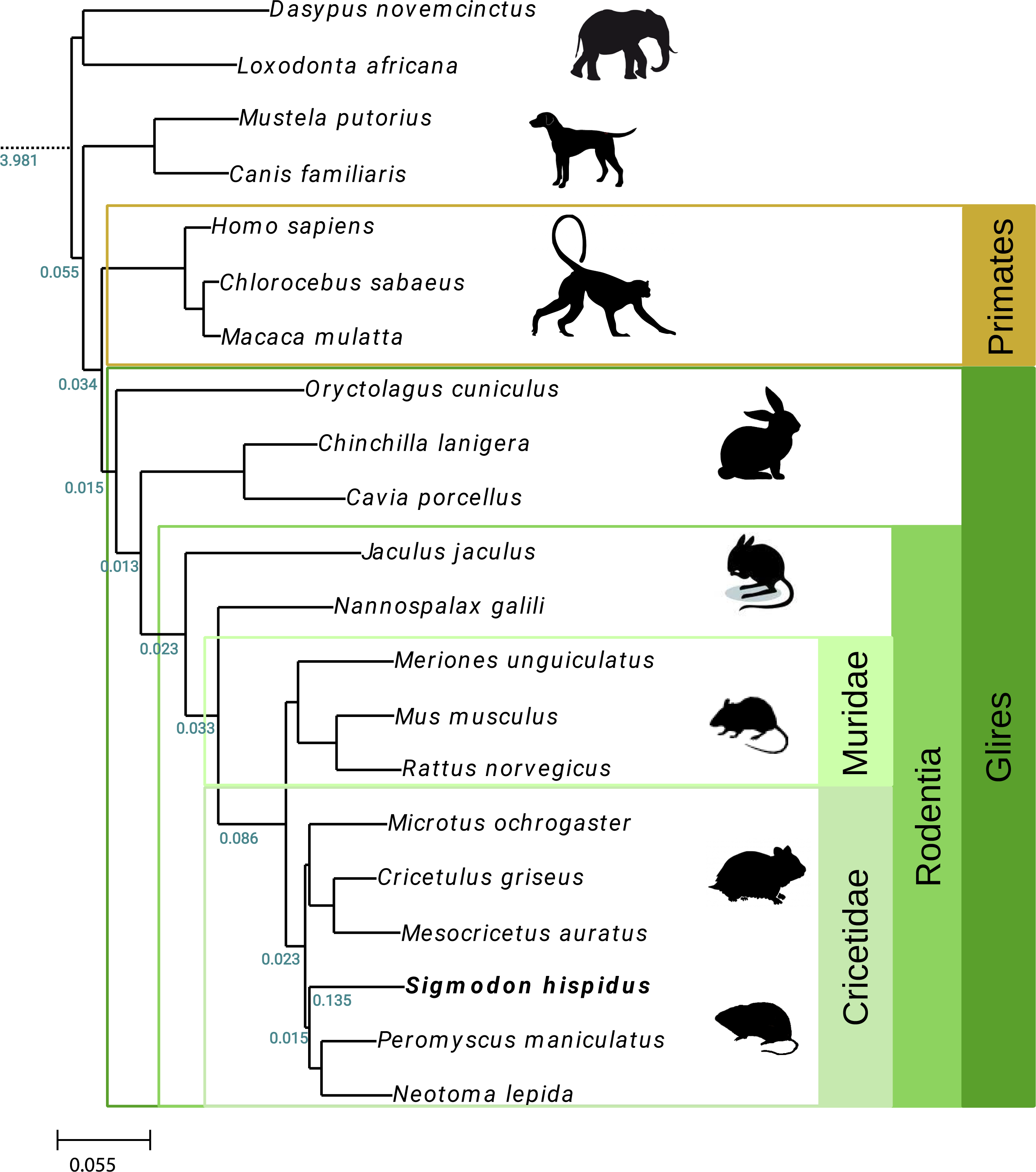
Species tree obtained from phylome analysis of multiple vertebrate species. A phylogenetic tree displays evolutionary relatedness among multiple vertebrate, mammalian and rodent species as labeled (*right,* species phyla). Support for all nodes was at 100% using the rapid bootstrap approach as implemented in RaxML (Stamatakis 2014). Duplication values were calculated after removing large species-specific expansions. *Left, numbers in green*, numbers of duplications per gene per branch; *bottom left, key,* reference rate of duplication as indicated.

To evaluate this possibility further, we obtained divergence times of interest (Pacifici 2013) from timetree.org (Hedges et al. 2015). We then divided the number of duplications per gene by the amount of time estimated to separate each species from the common ancestor (**Fig. 3**). Even after applying this normalization, we still estimated that the human genome harbors roughly half the number of gene duplications compared with the rodents (i.e. cotton rat, 0.013; mouse, 0.015; and human, 0.0054). This normalized result again suggests higher rates of gene duplication events in rodents (Thybert et al. 2018).

Phylomes provide a view of what happens to each gene over evolutionary time, and yield redundant data about gene duplications. To assess how many species-specific expansions are present, we clustered them using UPGMA (Ponce de Leon Senti 2017). We applied the condition that if at least 50% of proteins that have expanded within a cluster were found to overlap between clusters, then they were combined into larger clusters. We identified 206 such clusters containing five or more members each, with the largest cluster containing 269 proteins, including proteins of unknown function harboring the conserved domain “MTH889-like” (also of unknown function).

### Gains and losses of immune response gene homologs

Gains and losses of genetic orthologs and paralogs have been described among members of various gene families when studied across species. Such changes in gene counts particularly are anticipated when comparisons are made between species that diverged more than 20 million years ago (Lilue et al. 2013)(Nakaya et al. 2017). We used FatiGO (Al-Shahrour, Diaz-Uriarte and Dopazo 2004) to identify enriched gene ontology (GO) terms categorizing genes that are duplicated in the focus species. For cotton rat duplicons, a total of 199 GO terms were enriched, including several terms related to the immune system such as antigen binding, immunoglobulin production, immune response, complement activation and cellular response to interferon-gamma (**Supp. Table S5A**). We also identified GO terms for genes that are duplicated in mouse: 92 GO terms were found, of which several were related to the immune system (**Supp. Table S5B**). Similarly, 50 enriched GO terms for genes duplicated in human were found; again, several were related to immune functions (**Supp. Table S5C**).

We searched phylome trees for genes duplicated both in cotton rat and one of the other two focus species examined, i.e. duplicated either in cotton rat and human, or in cotton rat and mouse. This analysis identified 263 duplicated genes in both cotton rat and human, and 497 duplicated genes in both cotton rat and mouse. GO enrichment analysis of these duplicated genes showed several enriched ontology terms (9 in cotton rat and human, and 58 in cotton rat and mouse), which again included several immune system-related genes (**Table 4**).

**Table 4.** Comparisons of ontologies and biological processes involved in gene duplication events across three focus species.

| GO_TERM representative | cotton rat | mouse | human |
| --- | --- | --- | --- |
| antigen processing and presentation of peptide or polysaccharide antigen via MHC class II | No | Yes | No |
| complement activation | Yes | No | Yes |
| endocrine pancreas development | Yes | No | No |
| epoxygenase P450 pathway | No | No | Yes |
| establishment or maintenance of epithelial cell apical/basal polarity | No | No | Yes |
| flavonoid biosynthesis | No | No | Yes |
| glycolytic process | Yes | No | No |
| macrophage apoptotic process | No | No | Yes |
| mitotic prometaphase | Yes | No | No |
| negative regulation of cell differentiation | Yes | Yes | Yes |
| negative regulation of retinoic acid receptor signaling pathway | No | No | Yes |
| protein localization to adherens junction | No | No | Yes |
| receptor-mediated endocytosis | Yes | No | No |
| response to stimulus | No | No | Yes |
| sensory perception of smell | Yes | Yes | Yes |
| Sulfation | Yes | No | No |
| zonula adherens assembly | No | No | Yes |

Focusing further on particular genes whose involvement in human or mouse immune responses was established experimentally, we evaluated their gains or losses in the phylome trees (**Fig. 3**; **Supp. Fig. S6**). We tabulated 1,378 human immune response genes, of which 1,318 are found in phylomeDB, a public repository of gene phylogenies (Fuentes et al. 2022). Of these, 1,287 had orthologs in at least one other focus species, including 949 proteins with a cotton rat ortholog, and 1,024 with a mouse ortholog.

We carefully examined 352 immune-related genes belonging to 17 gene families (**Supp. Fig. S7**). Applying the neighbor-joining method to generate phylogenetic trees for each of these families (**Supp. Fig. S6**), we identified significant gains and losses of gene orthologs in all three focus species. Complicating this analysis, we observed that many genes annotated with the same name in both the mouse and human reference genomes were not direct orthologs. For example, human *MX1* and *MX2* paralogs share the same genetic ancestor, but they are distinct from both the mouse and cotton rat *Mx1* and *Mx2* paralogs that have a distinct molecular ancestor (**Supp. Fig. S6**). Similarly, the human *IFITM1*, *IFITM2* and *IFITM3* paralogs originated from a human-specific duplication event, but they are only distantly related to the mouse *Ifitm1*, *Ifitm2*, *Ifitm3*, *Ifitm6* and *Ifitm7* paralogs (**Supp. Figs. S6, S7**). This result documents that despite similar name assignments, gene paralogs can diverge markedly, which results in confusing or misleading gene nomenclature. In the cotton rat genome, we identified 11 *Ifitm* gene family members.

In tabulating losses of immune genes, we counted 63 absent from primates, 24 genes lost from humans, and 103 genes lost from cotton rat. We focused analysis on several functionally important genes absent from the cotton rat genome. For example, while human and mouse have one and two copies of *IFIT3* (human) and *Ifit3* and *Ifit3b* (mouse), respectively, cotton rat lacks an orthologous gene. Similarly, while both human and mouse genomes encode for *Apol6*, *Ccl1*, *Ang*, *Gsdmc*, *Sp100* and *Sp140*, each of these also is absent from cotton rat. An important minor histocompatibility antigen (MHA) locus that is present in mouse and human, i.e. the *Raet1/H60* locus (Gilfillan et al. 2002; Chalupny et al. 2003; Jung et al. 2012), is absent from the cotton rat genome assembly (**Supp. Figure S8**).

### Retrotransposons and other repetitive elements

Mobile genetic elements comprise approximately half of mammalian genomes, including those of all rodent species sequenced to date. They also play important roles in shaping genomes and contributing to speciation through evolution, and have been implicated in the etiology of diseases in human and other species (Cordaux and Batzer 2009; Hancks and Kazazian 2016; Chuang et al. 2021).

We used RepeatModeler to identify 1,534 distinct repetitive element families in the cotton rat genome (Smit 2008; Flynn et al. 2020). Resulting consensus sequences were used to define families in RepeatMasker, to count family members and to investigate divergence of individual family members from each family’s consensus. We identified 3.9 million repetitive elements in the cotton rat genome (**Supp. Table S6**). The ∼1.4 million SINE elements in the cotton rat genome outnumber ∼ 1.2 million SINEs elements in *M. musculus* genome (GRCm38; UCSC Mm10 mouse genome assembly). Similarly, ∼670,000 LTR elements in cotton rat outnumber ∼618,000 LTRs elements in mouse. However, ∼500,000 LINEs in the cotton rat are outnumbered by the >590,000 L1s in mouse genome (Mouse Genome Sequencing et al. 2002)(Richardson et al. 2015). While 19.9% and 17.5% of mouse and human genomic DNA is comprised of L1 sequences, respectively, only 15.5% of cotton rat genomic DNA is made up of L1. We count a total of 705 million bp comprising interspersed repetitive elements in cotton rat, compared with 946 million bp in mouse. The total fraction of the cotton rat genome comprised of repetitive elements is 38.8%, compared with 43.5% in mouse (mm10 assembly) (**Supp. Table S6**).

A large majority of repetitive elements identified in cotton rat have diverged more than 2% from their consensus sequences (**Fig. 5A**); these are considered to be ancient, inactive elements. By contrast, we identified only 16,384 repetitive elements with less than 2% sequence divergence from the consensus sequences, which represent young, active (recently retrotransposed) repeat family members. Of these, 59% (n=9,676) are SINE elements, and 36% (n=5,919) are LINE (L1) elements (**Fig. 5B**). We observed 740 LTR elements (4.5%), but none of these LTR elements are full-length LTR transposons.

**Figure 5.**
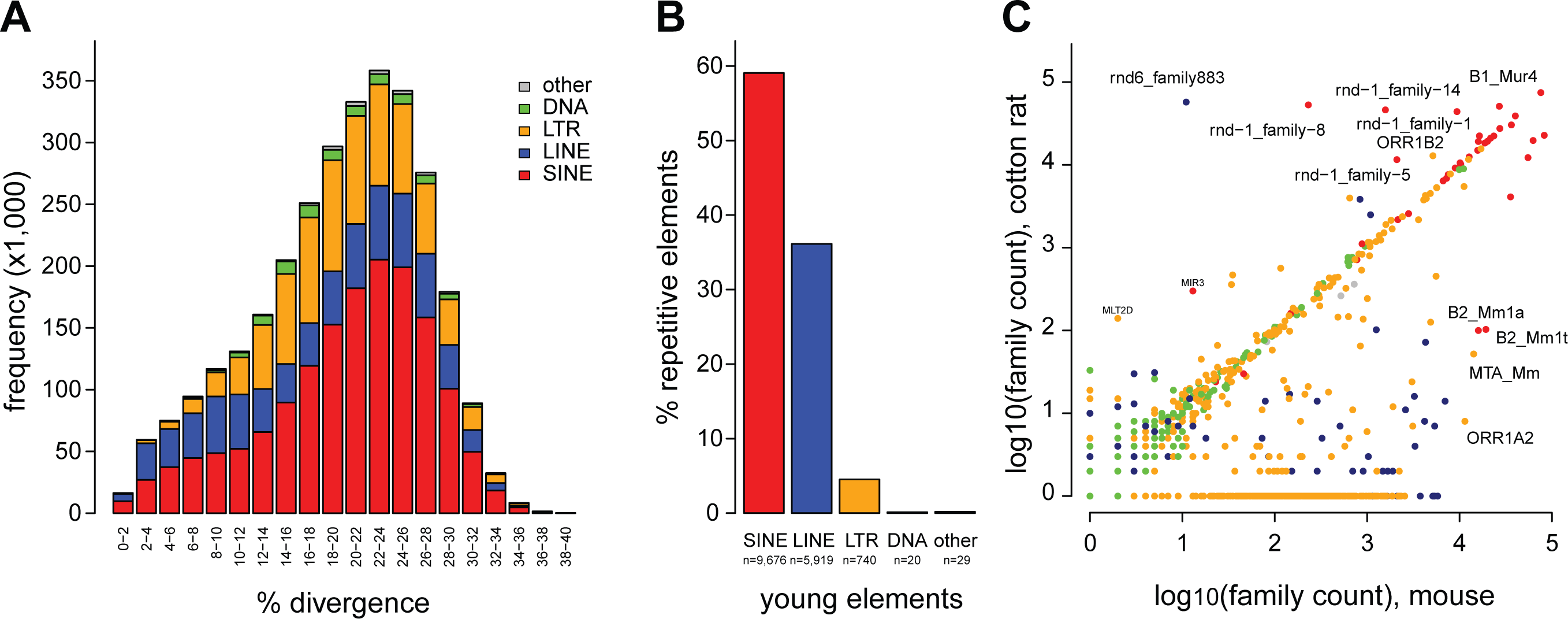
Repetitive elements in cotton rat genome. (A) Stacked bar graph depicts (*y-axis*) counts of (*key, upper right*) families of repetitive elements which (*x-axis*) diverge to various extents from consensus sequences as provided to RepeatMasker. *Key, colors*: *gray*, other; *green*, DNA repetitive elements; *orange*, LTR retroelements; *blue*, LINE retrotransposons; *red*, SINE retrotransposons. (B) Fraction of “young” repetitive elements in each repeat class. We identified 16,384 young repetitive elements in the cotton rat genome assembly, each with less than 2% divergence from respective consensus element sequences. *Key, colors*, as in panel A. (C) Scatterplot compares frequencies of related repetitive element families in genomes of (*y-axis*) cotton rat compared with (*x-axis*) mouse. Families of repetitive elements (*dots*) above the diagonal line (*line of identity*) are relatively enriched in the cotton rat genome, and include (*dark blue*) LINE (rnd6_family883); (*red*) SINE (rnd-1_family-8; rnd-1_family-14; and rnd-1_family-5); and (*orange*) LTR (MLT2D) families. For counting of individual family members, coverage was required to be >80%. Counts are log_10_-transformed. *Key, colors*, as in panel A.

We compared counts of repetitive element family members in cotton rat with those in *M. musculus* (**Fig. 5C**). As expected, most repeat classes included comparable numbers of elements in both genomes, most of which are presumably older, ancestral elements. We identified several distinct families of SINE and L1 elements that are markedly enriched in the cotton rat genome compared with the mouse, suggesting their relatively more recent mobilization after divergence of these species (**Fig. 5C**).

Members of one cotton rat SINE family (i.e. rnd-1-family-8#SINE/B2) represented 92% of all young SINE elements identified (**Supp. Fig. S9**). We refer to this rnd-1-family-8#SINE/B2 consensus as the cotton rat SINE B2 family (**Supp. Fig. S10**). This cotton rat SINE family is closest to the mouse SINE B3 family, but still exhibits 9.6% divergence compared to the mouse family.

Among the young LINE elements identified in cotton rat, we initially detected six L1 families each with more than 100 elements and having less than 2% divergence from the consensus sequences. After closely examining the consensus sequences of these six families, we noticed that they all closely overlapped, as five of the six families lacked the 5’ sequences present in the sixth family. Thus members of these five families were reclassified as 5’ truncation elements and members of the same L1 family, i.e. rnd-6_family-883#LINE/L1. We refer to this L1 family as L1sh (*S. hispidus*). The L1sh consensus sequence is closest to the mouse L1 Lx3_Mus family, but still is 20.1% divergent from the latter.

We examined the predicted protein-coding sequences in consensus L1sh elements. All mammalian L1 retrotransposons identified to date contain two open reading frames (ORF1 and ORF2), with ORF2 encoding endonuclease and reverse transcriptase activities. As expected, two ORFs were identified in L1sh. (**Supp. Fig. S11**). Comparison of L1sh ORF2 amino acid sequences revealed 74.6% identity with rat L1 ORF2 and 73.1% identity with mouse L1 ORF2. L1sh ORF1 has 67% identity to rat L1 ORF1 and 67% identity to mouse L1 ORF1. In general, mammalian L1 ORF2 proteins typically are more highly conserved across species than are L1 ORF1 proteins (Wagstaff et al. 2011); this trend holds in cotton rat.

To examine evolution of L1 retrotransposons across rodent and other mammalian species, we compared ORF2 proteins across ten vertebrate species by creating a phylogenetic tree (**Supp. Fig. S12**). The results are consistent with recent reports about evolutionary similarities in Sigmodon L1 elements (Yang, Scott and Wichman 2019).

## DISCUSSION

The cotton rat (*S. hispidus*), a rodent native to the Americas, has emerged as an animal model useful in the study of the pathogenesis of human respiratory viruses and in the development of vaccines and antiviral therapeutic agents. Cotton rats have innate susceptibility to a variety of human pathogens, although the genetic basis for this susceptibility has remained unknown. Here we report a new, high-quality genome assembly, based first upon our linked-read and PCR-free WGS data. By incorporating data from chromosome paints derived from *M. musculus* and comparative fluorescence microscopy analysis of metaphase spreads of several rodent species, i.e. *S. hispidus*, *M. musculus* and *P. leucopus*, we were able to make chromosome-level assignments in the assembly. While long stretches of chromosomal sequences display synteny with other rodent species, we also identified numerous examples of genetic innovation in gene families, supported by phylogeny analysis. A large majority of RNA-seq reads from pooled cotton rat tissues and independently from infected animals aligned well to annotated genes and even revealed alternative splicing isoforms, suggesting broad utility of this new genome assembly as a valuable reference in future studies.

This high-quality, highly contiguous genome assembly was generated by combining 10x linked-read and PCR-free WGS data, i.e. from short sequencing reads only (**Table 1**). We took advantage of two features of our experimental design: a high physical coverage of the genome was attained without PCR amplification-induced artifacts; and long-distance genomic connectivity between barcoded, linked reads was reached because they were derived from the same physical molecules (frequently >50-100 kb), thus contributing to the assembly (Ott et al. 2018). However, despite having a high scaffold N50 length, our intermediate assembly still was characterized by a relatively high number of small gaps. We patched them by generating complementary assemblies made from the PCR-free paired end libraries, followed by a second step where the remaining gaps were closed using Compass, a gap-filling software tool (https://github.com/nygenome/compass).

Comparison between our cotton rat genome assembly and another independent genome assembly that was released recently, i.e. SigHis_v1_BIUU, confirms the overall size and general composition of the cotton rat genome (**Table 1**). However, a limitation of the BIUU assembly is highlighted by its much higher counts of scaffolds, most of which are single-contig scaffolds reflecting its reduced contiguity and increased fragmentation overall. We observed examples of such fragmentation disrupting gene cluster structures in this recent BIUU assembly, which by contrast were resolved in our new genome assembly (**Fig. 2**; data not shown).

We took advantage of a relatively high degree of orthology between the *S. hispidus* genome and other rodents including the well-characterized *M. musculus* genome, by analyzing metaphase spreads from the former using chromosome-specific paints derived from the latter. This analysis facilitated assignment of *S. hispidus* genomic sequences into chromosomes on the basis of their lengths, banding patterns and synteny, resulting for the first time in a chromosome-level cotton rat genome assembly.

Genes in the new cotton rat reference genome were identified and annotated by making *ab initio* gene predictions, adding alignments of RNA sequencing data, and conducting cross-species comparisons. We estimated 24,878 protein-coding genes based on *ab initio* predictions (**Table 2**). This count is on par with other mammalian and rodent species. Of these, we detected expression of transcripts for 21,417 (96%) in RNA-seq libraries pooled from multiple cotton rat tissues (**Table 3**). Detailed examination of transcripts expressed from particular genes in several conserved immune gene families, upon alignment of RNA-seq reads against the annotated genome assembly, corroborated their predicted exonic structures. We identified multiple splicing isoforms for members of the *Ifit1*, *Cxcl* and *Gbp7* gene families, clustered on cotton rat Chrs. 2, 1 and 16, respectively. These data were confirmed further by alignment of independent RNA-seq libraries generated from cotton rat individuals exposed to RSV infection (Strickland et al. 2022).

Phylome analysis facilitated investigation of the evolutionary relatedness amongst members of gene families conserved among the cotton rat, mouse, human and other species. This revealed numerous genomic innovations manifested as gene family gains and losses distinguishing cotton rat from other rodents. The finding that cotton rat has undergone a more pervasive loss of immune-related genes compared to mouse and humans may have implications about its increased susceptibility to infection by viruses and other pathogens. However, we acknowledge that some of these apparent gene losses may be attributed in part to technical difficulties in assembling short-read sequencing data de novo. In particular, numerous immune genes (e.g. in the major histocompatibility complex, MHC and immunoglobulin gene loci) are encoded in repetitive gene clusters that are particularly difficult to resolve from short reads (Lilue et al. 2018).

Functional studies have highlighted both some differences between mouse and cotton rat immune responses and concomitant similarities between the latter and humans. In contrast to the mouse but similar to humans, cotton rats have columnar epithelial cells in their respiratory tract, which are the target of infection by viruses such as RSV and influenza virus (Grieves et al. 2015; Green et al. 2021). Toll-like receptor 9 molecules are expressed at much lower levels in cotton rat or human lymphoid tissues than in the mouse (Kim and Niewiesk 2014). Human *TLR9* agonists also stimulate cotton rat cells better than agonists developed for mouse *Tlr9* (Kim and Niewiesk 2013; Kim and Niewiesk 2014). As in human macrophages, cotton rat macrophages produce little nitric oxide, implying that nitric oxide is mainly used as a signal transduction molecule in both species, and not as an anti-bacterial effector as it is in mice (Carsillo et al. 2009). Another similarity between humans and cotton rats lies in the constitutive expression of heat shock protein 70 (*HSP70*) in tissues, which is not the case in mice (Niewiesk, unpublished). We anticipate that the availability of a new reference genome will facilitate further genetic studies on the basis for these and other functional properties in cotton rat.

More than 800 cotton rat genes’ coding sequences previously were cataloged in the National Center for Biotechnology Information (NCBI) database. Most of these are related to the immune response. We compared individual orthologs in cotton rat, mouse, human and other species, in an effort to decipher differences between their susceptibility and immune responses to various pathogens. Overall, we found that the cotton rat genes are more closely related to hamster genes than to mouse and rat genes (data not shown). This evolutionary relationship was illustrated by orthologs of *CD1*, a restriction element of natural killer (NK) cells (Fichtner et al. 2015). The observed differences in homology are consistent with the split between the Muridae (mouse, rat) and Cricetidae (hamster, cotton rat) families 23.3 to 24.9 million years ago, whereas the split between the Arvicolinae-Cricetinae (hamster) and Sigmodontineae-Netominae (cotton rat) occurred 18.7–19.6 million years ago (Steppan, Adkins and Anderson 2004). However, all of these rodent genes are more closely related to each other than to human genes.

*CD150* (a lymphocyte activation molecule) is another annotated gene with important functions, as it encodes a receptor for measles virus (MV) that is functional in human and cotton rat, whereas the mouse ortholog is not functional. Key differences in amino acids at positions 60, 61 and 63 may explain these interspecific differences in MV receptor function (Carsillo et al. 2014). In contrast, no such differences could be identified in the orthologous RSV receptor protein CX3CR1 encoded in cotton rats, mice and humans (Green et al. 2021). In this case, unidentified downstream factors may influence or mediate differential susceptibility to RSV infection. Like humans, cotton rats express a functional set of Mx proteins encoding the antiviral proteins Mx1 and Mx2 (Pletneva et al. 2006). These proteins severely reduce infection of the cotton rat lung tissue by influenza virus. In contrast, most mouse strains lack functional Mx proteins, and therefore rely on an incomplete innate type I interferon response.

The draft reference genome of cotton rat defines the structures of several dozen gene candidates involved in virus susceptibility or immunity, and therefore will facilitate future research on many aspects of viral replication and immunity in cotton rats. We have documented a significant number of losses of particular immune-related genes, absent from the cotton rat genome assembly. Some genes that are conserved between human and mouse but are confirmed to be absent from cotton rat include *Ccl1*, *Ang* gene family members, *Ifit3*, *Rarres2*, *Nlrp2*, *Gsdmc*, *Reat1e Sp100* and *Sp140*. Other genes are lost from both human and cotton rat but present in mouse (although mouse and cotton rat are evolutionarily closer to each other), including AIM2-like receptor members (*Pydc3* and *Pydc4*), *H60b*, *Ifitm6*, *Irgb10*, *Irga6* and *Apol6*. The loss of these genes from the cotton rat genome may help explain the host’s exquisite susceptibility to human viruses and other pathogens. Further characterization of the potentially protective roles of these genes in immunity is warranted.

We acknowledge several ways by which this new cotton rat genome assembly could be improved with additional studies. First, although linked-read sequences are derived from long physical DNA molecules and therefore facilitate analysis of long-range connectivity spanning repetitive genomic elements, they nevertheless are discontinuous. Despite improvements in linked-read library preparation methods, identical barcodes still frequently label multiple independent genomic DNA segments, thereby introducing potential artifacts in assemblies. Other methods such as continuous long-read sequencing, optical mapping and Hi-C methods have been optimized recently. Use of the latter methods is likely to improve connectivity spanning across widespread repetitive elements that is required for high-quality genome assemblies (Rhie et al. 2021). Second, the individuals sequenced here were highly inbred, so extensive genomic homozygosity would be expected. Genomic studies in diverse, wild-caught, outbred cotton rats would help define population-level allelic variation frequencies and elucidate the population size. Third, while some chromosome-level scaffolds likely represented second haplotypes in genomic regions where heterozygosity persisted, we did not explicitly address allelic variation or diploidy here, nor did we conduct phasing. Identification of additional haplotypes will be facilitated by analysis of diverse cotton rat individuals. Inclusion of the additional sequencing data supporting the recently released, independent BIUU assembly would likely add such haplotype information. In adddtion, the resulting “combined” genome assembly would be improved from increased depth of sequencing coverage. And finally, a mean of only 1.17 transcripts was detected per annotated gene. Deeper RNA-seq across a full complement of tissues and diverse experimental and developmental conditions would improve quantification of tissue-specific expression of most annotated genes and identification of novel transcripts. In addition, many more examples of alternative splicing and diverse transcript isoforms would be identified.

This first high-quality, chromosome-level assembly of the cotton rat genome resulted from a combination of PCR-free whole genome sequencing, linked-read sequencing, and chromosome paint data, along with RNA-seq data. The development of this reference genome has allowed us to assess evolutionary relatedness between genomes of the cotton rat and other rodent species, some of which also are used frequently in research studies. This analysis has confirmed the cotton rat as a new world rodent, in contrast to mice and rats. These findings are extended further by chromosome assignments (karyotype 2n=52) which demonstrate significant similarity and synteny to other new world rodents, and extensive rearrangements when compared to the mouse. Cotton rats also differed from mice in the relative frequencies of SINE, LINE and LTR elements observed. We identified several families of transposable elements unique to *S. hispidus* that actively contributed to its distinctive genomic structure, including L1sh, distinguishing the cotton rat from other evolutionarily related rodent species.

## CONCLUSIONS

To assemble and annotate a high-quality, reference assembly of the cotton rat genome, we combined multiple lines of evidence including PCR-free whole genome sequencing, linked-read sequencing, chromosome paint data, and RNA-seq data. This genome reference is extended further by chromosome assignments (karyotype 2n=52) which demonstrate significant similarity and synteny to other new world rodents, and extensive rearrangements when compared to the mouse. Several unique families of transposable elements were identified in *S. hispidus*, which actively contributed to its distinctive genomic structure. They included L1sh, distinguishing the cotton rat from other evolutionarily related rodent species. We anticipate that this annotation and characterization of a chromosome-level cotton rat genome assembly will serve as a valuable resource, and will facilitate and accelerate ongoing investigations into its host defenses against viral and other pathogens, genome biology, and mammalian evolution.

## MATERIALS AND METHODS

### Animals

Inbred cotton rats (*S. hispidus*) that were between 4 and 8 weeks of age and free of specified pathogens (as specified by the breeder) were purchased from Envigo, Inc. (Indianapolis, IN). They were maintained in a barrier system in accordance with a protocol approved by the Ohio State University Institutional Animal Care and Use Committee. Environmental conditions were maintained at 20±2° C and 30-70% relative humidity with a 12 h light cycle. Euthanasia was performed via CO_2_ inhalation.

### Genomic DNA extraction, library preparation and sequencing

To obtain high molecular weight DNA for linked-read sequencing and WGS library preparation, DNA was extracted from an individual male cotton rat’s ear pinna tissue, using MagAttract HMW DNA kit (Qiagen). The isolation protocol including RNase treatment followed the manufacturer’s recommendations. DNA quality and concentration and the DNA integrity number were measured using a Nanodrop spectrophotometer, Qubit fluorimenter and an Agilent TapeStation, respectively.

To generate PCR-free libraries, we fragmented genomic DNA using a Covaris S2 sonicator, resulting in DNA fragments of various size distributions. We optimized shearing to yield median fragment lengths of 350 and of 550 nt in two independent aliquots. An Illumina TruSeq DNA Sample Prep kit was used to add adapter and sample barcode sequences to the resulting genomic DNA fragments, following the manufacturer’s protocol. Library quality was assessed using an Agilent TapeStation, and concentrations were determined using a Qubit fluorimeter. Sequencing was performed on an Illumina HiSeq2500, resulting in 2×150 bp paired-end reads with indexes incorporated for each library.

A linked-read genomic DNA library was prepared from approximately 1.25 ng high molecular weight genomic DNA as template, using the 10x Genomics Chromium genome library and gel bead kit (10x Genomics) to create Gel Bead-In-EMulsions (GEMs), following the manufacturer’s protocol. Isothermal incubation of the GEMs produced DNA fragments barcoded with10x Genomics linked-read indexes. Additional sequencing primers and sample indexes were added by end repair, A-tailing and adaptor ligation. After amplification, the barcoded library was size selected. The resulting library structure and concentration were assayed by quantitative PCR (KAPA Biosystems). Sequencing was conducted on an Illumina HiSeq2500, yielding 2×150 bp paired-end reads along with the sample and 10x linked-read molecular indexes.

### Bioinformatic analysis of sequencing data

In an initial pre-processing step, reads from the two PCR-free libraries were screened for adapter sequences and low-quality bases (Q<10) and trimmed using Cutadapt 1.8.1 (Martin 2011). After this step, read pairs containing a single read shorter than 50bp were discarded. To filter out spiked-in sequences, remaining reads were mapped against the PhiX reference sequence using GEM mapper (edit distance <= 10%) (Marco-Sola et al. 2012). Finally, processed PCR-free reads were error-corrected using Lighter v1.1.1 (k=21) (Song, Florea and Langmead 2014).

10x Chromium linked-read data also were error-corrected, using bloom-filters generated from the PCR-free data which were characterized by lower error rates. Barcodes and an additional seven nucleotides were clipped off from each Read 1 prior to error-correction. They were re-included subsequently, to ensure valid inputs for the assembler.

### Genome assembly

Error-corrected 10x linked reads were used as inputs into Supernova v2.0.1 (Weisenfeld et al. 2017). A pseudo-haploid representation of the assembly was generated using the subcommand *mkoutput*. The assembly was further scaffolded using ARKS v1.0.2 (Coombe et al. 2018).

Processed PCR-free data was used to produce nine contig assemblies using ABySS 2.0.2 (Jackman et al. 2017), exploring different *K*-mer sizes (i.e., 37, 47, 57, 67, 77, 87, 97, 107 and 117). Flanks of decreasing lengths (starting at 1kb and ranging down to 100bp, in decrements of 100bp) around each gap in the Supernova assembly were searched for in these assemblies. When both flanks mapped unambiguously to the same contig and in the correct order and orientation, using GEM mapper (Marco-Sola et al. 2012), the sequence between the outermost mapping coordinates was extracted and used to patch the gap, giving priority to sequences originating from assemblies of larger *K*-mer size. Remaining gaps were filled using Compass (https://github.com/nygenome/compass), exploring the same *K*-mer sizes listed above. Finally, all scaffolds shorter than 200 kb were searched for in the assembly, using MegaBLAST (Zhang et al. 2000). Scaffolds that fully aligned to a larger scaffold (coverage = 100%, identity >=99%) were considered redundant and therefore removed.

Gene completeness was evaluated using CEGMA 2.5 (Parra, Bradnam and Korf 2007) with a default 248 core eukaryotic gene set, and using BUSCO 5.4.0 (Simao et al. 2015) with the *mammalia_ odb10* gene set. In another check of completeness, PCR-free data were mapped in paired-end mode against the final assembly with BWA-MEM v0.7.17 (Li 2013). Corresponding mapping statistics were computed using CollectMultipleMetrics from the Picard toolkit v2.16.0 (https://broadinstitute.github.io/picard).

### Assigning pseudo-chromosomes

Preparations of cotton rat chromosomes were made from embryo fibroblast cells that were isolated from six 12-14 day old cotton rat embryos using a mouse embryonic fibroblast isolation kit (Pierce). Metaphase chromosome preparations and cross-species chromosome painting were performed using laboratory mouse chromosome probes as described previously (Yang, O’Brien and Ferguson-Smith 2000) (**Supp. Fig. S1**). Paint probes specific for *S. hispidus* Chrs. 1, 2, 3, 8, 9, 11, 22, 23, 24 and 25, generated from flow-sorted mouse chromosomes, also were hybridized onto *O. torridus* metaphase chromosomes to evaluate concordance between chromosomes of *S. hispidu*s and *O. torridus*.

Pseudo-chromosome models were constructed using Syn2Chr (https://github.com/igcbioinformatics/Syn2Chr/), based on synteny between cotton rat and the mouse reference genome GRCm38. Ambiguous genome structures were corrected based on an updated *Peromyscus leucopus* genome assembly (i.e. UCI_PerLeu_2.1, https://www.ncbi.nlm.nih.gov/assembly/GCF_004664715.2) and on the *O. torridus* genome (i.e. mOncTor1.1, GCA_90399543, https://www.ncbi.nlm.nih.gov/assembly/GCF_903995425.1).

### RNA extraction and sequencing

Tissues included for analysis by RNA sequencing (RNA-Seq) were harvested freshly from euthanized male cotton rats. RNA was extracted from brain, heart, intestine, kidney, liver, lung, mediastinal lymph node, muscle, Peyer’s patch, spleen, and thymus of a healthy, naive adult cotton rat male. Independently, a second cotton rat male was injected intraperitoneally with 100 μg of house dust mite (HDM) adsorbed to AdjuPhos (Brenntag) in a 1:1 v/v ratio (Green et al. 2018). Eight days later, it was challenged intranasally with 100 μg of HDM in a 100 μl volume. The cotton rat was euthanized 4 days post-challenge, and RNA was extracted from lung, mediastinal lymph nodes, and spleen. RNA was isolated using the RNeasy Microarray Tissue kit (Qiagen).

RNAs from various tissues were pooled for each individual. RNA pool concentrations and quality were checked using a NanoDrop (Thermo Scientific) and Bioanalyzer (Agilent), respectively, and were determined sufficient for RNA sequencing. Messenger RNAs were enriched based on their poly(A) tails, reverse transcribed to cDNA and barcoded using indexed adapters to permit multiplexing of individual sample pools. The cDNA libraries were prepared and finished, and quality and concentration were measured, in the Ohio State University Comprehensive Cancer Center Genomics Shared Resource. Sequencing was carried out in a single lane in an Illumina HiSeq4000 instrument, generating 2×150 bp paired-end reads output in fastq file format.

### RNA-seq analysis

RNA-seq reads were aligned to the cotton rat genome assembly mSigHis_REL_1907.fa, harboring 26 chromosomes and more than 16,000 unplaced scaffolds, with annotation file mSigHis_REL_1907.gff3. We used STAR v2.5.3a (Dobin et al. 2013), using command lines including genomeDir genome --readFilesIn ../evidence/NaiveCR_1-11_RNAseq/1_11_S28_L007_P0001_R1.fastq.gz ; ../evidence/NaiveCR_1-11_RNAseq/1_11_S28_L007_P0001_R2.fastq.gz ; --readFilesCommand zcat ; --runThreadN 4 -- outFileNamePrefix star/Naive ; --outSAMstrandField intronMotif --outSAMtype BAM SortedByCoordinate ; --outSAMattrIHstart 0 ; --outFilterIntronMotifs RemoveNoncanonical ; -- outTmpDir $TMPDIR/Naïve.

For comparisons with RNA-seq data generated independently (Strickland et al. 2022), Illumina raw reads were downloaded from SRA database (https://www.ncbi.nlm.nih.gov/sra/?term=%20SRR23104992). Alignments against our cotton rat genome assembly were performed using STAR 2.79a with command lines: STAR --genomeDir ../Star2.79a -- readFilesIn <(zcat SRR23104982_1.fastq.gz) <(zcat SRR23104982_2.fastq.gz) --alignIntronMax 500000 --outSAMtype BAM SortedByCoordinate --outFileNamePrefix Inf1 --sjdbOverhang 99 –sjdbGTFfile ../mSigHis_REL_1907.gff3 --runThreadN 16.

### Gene annotation

Gene annotation of the cotton rat genome assembly were generated by combining transcript alignments, protein alignments and *ab initio* gene predictions. Transcript models were prepared using Cufflinks v2.2.1 (Trapnell et al. 2010). With the addition of 107 *Sigmodon* genes downloaded from NCBI, PASA assemblies were produced with PASA v2.3.3 (Haas et al. 2008). TransDecoder, part of the PASA package, was run on the PASA assemblies to detect coding regions in the transcripts. The complete human, rat and mouse proteomes were downloaded from Uniprot, and aligned to the genome using spaln (Gotoh 2008) (v2.2.2). *Ab initio* gene predictions were performed on the repeat masked cotton rat assembly using three different programs: GeneID v1.4 (Parra, Blanco and Guigo 2000), Augustus v3.2.3 (Stanke et al. 2006) and Genemark-ES v2.3e (Lomsadze, Burns and Borodovsky 2014), with and without incorporating evidence from RNAseq data. The gene predictors were run with the human trained parameters, except Genemark, which runs in a self-trained manner. All results were combined into consensus gene coding sequence (CDS) models using EvidenceModeler v1.1.1 (Haas et al. 2008). Additionally, untranslated regions (UTRs) and alternative splicing isoforms were updated through two rounds of PASA annotations. Functional annotations were assigned to proteins with Blast2go (Conesa et al. 2005). A Blastp (Altschul et al. 1990) search was run using the nr database and then Interproscan (Zdobnov and Apweiler 2001) was run to detect protein domains on annotated peptides. All resulting data were combined using Blast2go which produced the final functional annotation results.

Annotations of noncoding RNAs (ncRNAs) were produced by running the following steps. First, the program cmsearch v1.1 (Cui et al. 2016), part of Infernal (Nawrocki and Eddy 2013), was run against the RFAM (Nawrocki et al. 2015) database of RNA families (v12.0). In addition, tRNAscan-SE v1.23 (Lowe and Eddy 1997) was run to detect transfer RNA genes present in the genome assembly. To detect long-noncoding RNAs (lncRNAs), we selected those PASA-assemblies that had not been included into the annotation of protein-coding genes in order to evaluate expressed but untranslated transcripts. Particular PASA assemblies lacking protein-coding annotations that exceeded 200bp and whose length was not covered at > 80% by a small ncRNA were incorporated into the ncRNA annotation as lncRNAs. Resulting transcripts were clustered into genes using shared splice sites or significant sequence overlap as criteria for designation as the same gene.

### Phylome reconstruction

We constructed a phylome, i.e. a complete collection of phylogenetic trees depicting the relatedness of each gene across a set of evolutionarily distinct genomes. Cotton rat, mouse and human genomes were included in this analysis to identify gene gains and losses among these key focus species. Twenty additional species included 11 Rodentia species were investigated in each phylome. Phylomes were constructed using an automated pipeline (Huerta-Cepas and Gabaldon 2011; Fuentes et al. 2022). Briefly, for each predicted protein encoded by a particular genome, a Smith-Waterman search was performed against a proteome database (Smith and Waterman 1981). Results were filtered using an e-value cut-off <1E-5 and a continuous overlapping region of 0.5. Up to 150 homologous sequences for each protein were identified, which were then aligned using MUSCLE v3.8 (Edgar 2004), MAFFT v6.712b (Katoh et al. 2005), and kalign (Lassmann and Sonnhammer 2005). Alignments were performed in forward and reverse orientations and in each possible reading frame using the Head or Tail approach (Landan and Graur 2007). The resulting six alignments were combined with M-COFFEE (Wallace et al. 2006) and then trimmed with trimAl v1.3 (Capella-Gutierrez, Silla-Martinez and Gabaldon 2009), with a consistency score cutoff of 0.1667 and gap score cutoff of 0.9. Trees were reconstructed using the best-fitting evolutionary model. The selection of the model best fitting each alignment was performed as follows: a Neighbor Joining (NJ) tree was reconstructed as implemented in BioNJ (Gascuel 1997). The likelihood of each topology was computed, allowing branch-length optimization, using 7 different models (JTT, LG, WAG, Blosum62, MtREV, VT and Dayhoff), as implemented in PhyML v3.0; and then the model best fitting the data, as determined by the AIC criterion (Akaike 1973), was used to derive ML trees. Four rate categories were used, and invariant positions were inferred from the data. Branch support was computed using an approximate likelihood ratio test (aLRT), based on a chi-square distribution. Resulting trees and alignments are stored in phylomeDB (http://phylomedb.org), labeled with phylomeIDs 19 (cotton rat), 20 (mouse) and 21 (human). Trees were scanned using ETE v3.0 (Huerta-Cepas, Serra and Bork 2016).

### Species tree reconstruction

A total of 895 proteins were found encoded as single-copy genes in all 21 species. These proteins were used to reconstruct a species tree by concatenating the clean alignments produced during phylome reconstruction. The concatenated alignment contained 635,820 amino acid positions. RAxML HPC-PTHREADS-SSE3 version 8.2.4 (Stamatakis 2014) was then used to reconstruct a phylogenetic tree based on the PROTGAMMALG model. Branch support was calculated using the rapid bootstrap approach implemented in RAxML. Species trees followed the taxonomic classification as expected.

### Orthology and Paralogy determination

Trees were scanned for orthologs and paralogs using a species overlap algorithm as implemented in ETE v3.0 (Huerta-Cepas, Serra and Bork 2016). For each tree, nodes were annotated as speciation or duplication nodes, depending on whether there were common species at both sides of the node or not. When common species were present, a duplication node was annotated. In this case, sequences on either side of the node were considered as paralogs. If no common species were found, a speciation node was annotated. In such a case, the sequences were considered as orthologs.

### GO term assignment and enrichment analysis

GO terms for proteomes included in the phylomes were downloaded from phylomeDB. The GO terms were transferred between one-to-one and many-to-one orthologs to cotton rat genes. Enrichment of GO terms was calculated using a python adaptation of FatiGO (Al-Shahrour, Diaz-Uriarte and Dopazo 2004).

### Annotation of transposable elements

RepeatModeler version open-1.0.11 (Smit 2008) was used to detect repeat families in the cotton rat genome assembly (NYGC NIEWIESK1701.v2.fa), identifying 1534 repeat families. Resulting repetitive family consensus sequences were used as input for RepeatMasker version open-4.0.7 (Smit 2013) to count repeat elements and their divergence from consensus sequences. To count L1 elements, we used NCBI ORFfinder (Wheeler et al. 2003) to detect the open reading frames in cotton rat L1 LINE elements and determine the amino acid sequence of the L1 ORF2 reverse transcriptase. The amino acid sequence of the reverse transcriptase was compared with other vertebrate L1 reverse transcriptases using Clustal Omega (Madeira et al. 2019).

## DECLARATIONS

### Ethics approval

Housing, experimental treatment and euthanasia of cotton rats as described in this manuscript was reviewed and approved by the Institutional Animal Care and Use Committee at Ohio State University.

### Consent for publication

Not applicable

### Data availability

RNA-seq, WGS data and the genome assembly and annotations from this study have been deposited in the NCBI repository under BioProject PRJNA720389 with RNAseq data (SUB157345), WGS data (SUB9369558) and genome assembly. Annotated chromosome-level genome assembly and aligned RNA-seq data will be submitted as BED files for species-specific tracks at the UCSC browser. Genome annotations and metadata are available for visualization and searching via a genome browser and BLAST server at https://denovo.cnag.cat/cottonrat

### Availability of data and materials

RNA-seq, WGS data and the genome assembly and annotations from this study have been submitted to the NCBI repository under BioProject PRJNA720389 with RNAseq data (SUB157345), WGS data (SUB9369558) and genome assembly. In addition, all genome annotations and metadata are available for visualization and searching via a genome browser and BLAST server at https://denovo.cnag.cat/cottonrat.

### Authors’ contributions

**Conception of the work:** DES, SN; **design of the work:** JL, AC, JG-G, KA, TA, TG, BH, DES, SN; **acquisition, analysis or interpretation of data:** JL, AC, JG-G, FY, KA, GG, BLN, BF, UPC, SW, MM-H, TK, JCM, TA, TG, BH, DES, SN; **creation of new software used in the work:** none; **drafted or substantially revised manuscript:** JL, AC, JG-G, KA, TK, TA, TG, BH, DES, SN.

## Acknowledgments

Next-gen sequencing was performed in the Ohio State University Comprehensive Cancer Center (OSUCCC) Genomics Shared Resource, supported by NCI Cancer Center Support Grant P30CA016058.

## Authors’ information

N/A

