## Supplemental Information File for "A chromosome-level genome assembly of the Hispid cotton rat (*Sigmodon hispidus*), a model for human pathogenic virus infections"

**SUPPLEMENTARY INFORMATION**

**TABLE OF CONTENTS**

pg. nos.

**Supplementary Results** 3

**Supplementary Figures**

Supplementary Figure S1. Homologous chromosomes in cotton rat and mouse.

(a) Mouse chromosome paints identify homologous S. hispidus chromosomes. 4-9

(b) Chromosome homology map. 10

Supplementary Figure S2. Comparison of orthology and synteny between mouse and cotton rat chromosomes. 11-14

Supplementary Figure S3. Comparison of pseudo-chromosome assemblies of cotton rat (*S. hispidus*) and white-footed mouse (*P. leucopus*) reveals shared orthology with mouse. 15

Supplementary Figure S4. Comparison of orthology and synteny between *S. hispidus, O. torridus* and *P. maniculatus* chromosomes. 16

Supplementary Figure S5. Gene counts in cotton rat tissue pools. 17

Supplementary Figure S6. Phylogenetic trees of 17 immune-related gene families. 18-26

Supplementary Figure S7. Three-way comparison of 352 immune-related genes from 17 families. 27

Supplementary Figure S8. Loss of *Raet* / *MHA* locus from cotton rat. 28

Supplementary Figure S9. Counts of repetitive elements by family. 29

Supplementary Figure S10. Nucleotide sequence of consensus cotton rat SINE/B2 element. 30

Supplementary Figure S11. Predicted amino acid sequences of consensus cotton rat L1 ORF1 and ORF2. 31-32

Supplementary Figure S12. Phylogenetic tree characterizes evolution of vertebrate L1 ORF2s. 33

**Supplementary Tables**

Supplementary Table S1. Alignment features of bulk RNA-seq reads from cotton rat tissue pool P1. 34

Supplementary Table S2. Selected “immune response” (GO term) genes in cotton rat genome. 35

Supplementary Table S3. Immune response genes whose expression was not detected in cotton rat. 36

Supplementary Table S4. Confirmation and annotation of differential gene expression upon RSV infection in cotton rats. 37

Supplementary Table S5. GO terms of immune genes duplicated in the cotton rat genome. 38

Supplementary Table S6. Comparison of repetitive element counts in mouse vs. cotton rat genomes. 39

**Supplementary References** 40

### SUPPLEMENTAL RESULTS

A total of 155 gene losses (including 103 from house mouse, 89 from human and 37 shared; see **Supp. Fig. S7**) were manually inspected by identification of adjacent, syntenic genes in the mouse and human reference genomes. Among them, 58 (37.4%) were confirmed as being annotated by gene prediction pipeline, but most of these either had annotation problems or exceeded a divergence limit established for the phylome trees analysis. In addition, we confirmed the presence of 24 (15.5%) genes in our reference genome assembly, but they were not annotated. Another 47 (30.3%) were missed due to assembly gaps or unresolved long repeat regions (e.g. Defensin loci and H2/MHC locus). Interestingly, 26 gene losses were confirmed in cotton rat genome (**Supp. Fig. S7**), including genes playing key roles in mouse and human viral resistance, including *Oas1b* (Green et al. 2017), *Mndal* (Brunette et al. 2012), *CCL1* (Saito et al. 2017) and *Ifi6* (Qi et al. 2017).

*Copy number variation*

We focused our analysis on 352 immune-related genes belonging to 17 gene families, including the *Slfn* (Li et al. 2012), *Oas* (Green et al. 2017), *Nlrp* (Sarvestani and McAuley 2017), *Ifit* (Feng et al. 2018), *Ifitm* (Bailey et al. 2014), *Mx* (Verhelst et al. 2012) and *AIM2*-like receptor families (Nakaya et al. 2017), which encode antiviral factors in mouse and / or human. In addition, we evaluated members of the *Irg, Gbp*, *Cxcl* and *Cxcr* gene families, which have been shown to be significantly upregulated in *S. hispidus* lung transcriptome following respiratory syncytial virus infection (Rajagopala et al. 2018); the *ApoL*, *Ccl*, *Ang*, *Gsdm*, and *CD300* family members, which may have undergone gene losses in cotton rat, as well as the *Raet1/H60* locus. We didn't investigate *Sp100* genes further, because this gene family is not fully resolved in the current GRCm39 reference genome for house mouse (cf. https://www.ncbi.nlm.nih.gov/grc/mouse/issues?q=MG-3504).

**SUPPLEMENTAL FIGURES**

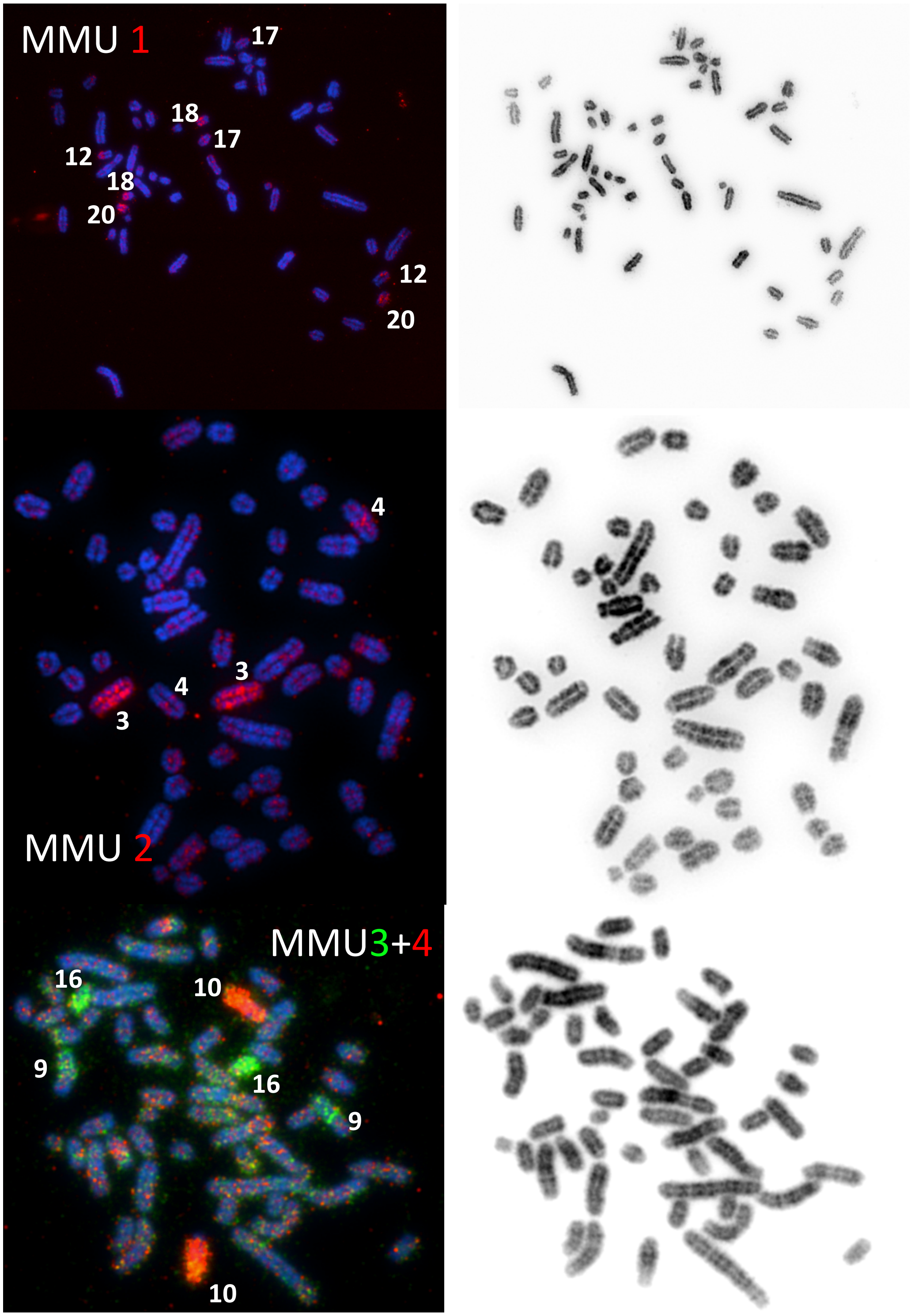

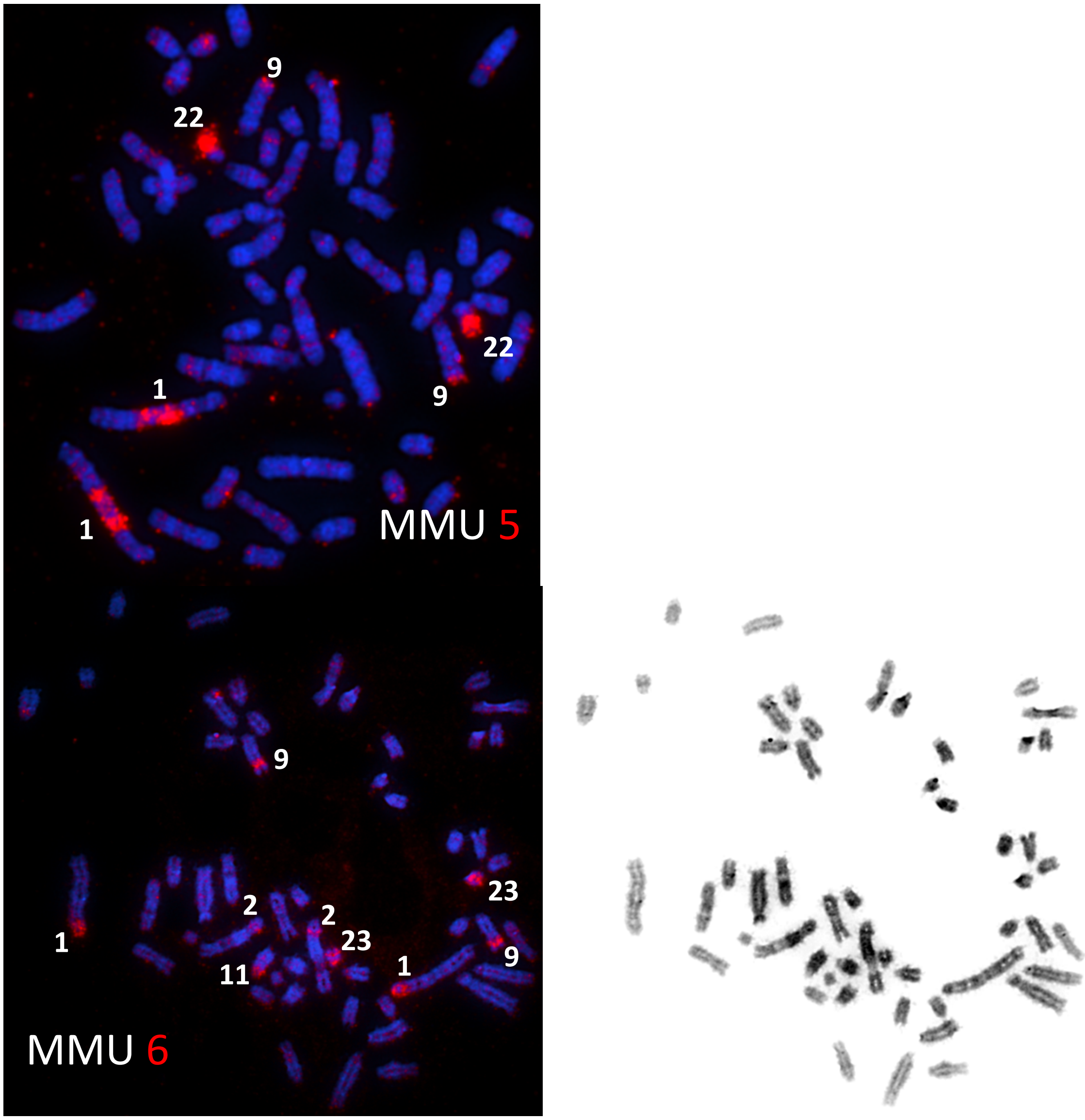
9
22
22
9
1
1
MMU 5
9
23
2
2
23
1
11
1
9
MMU 6

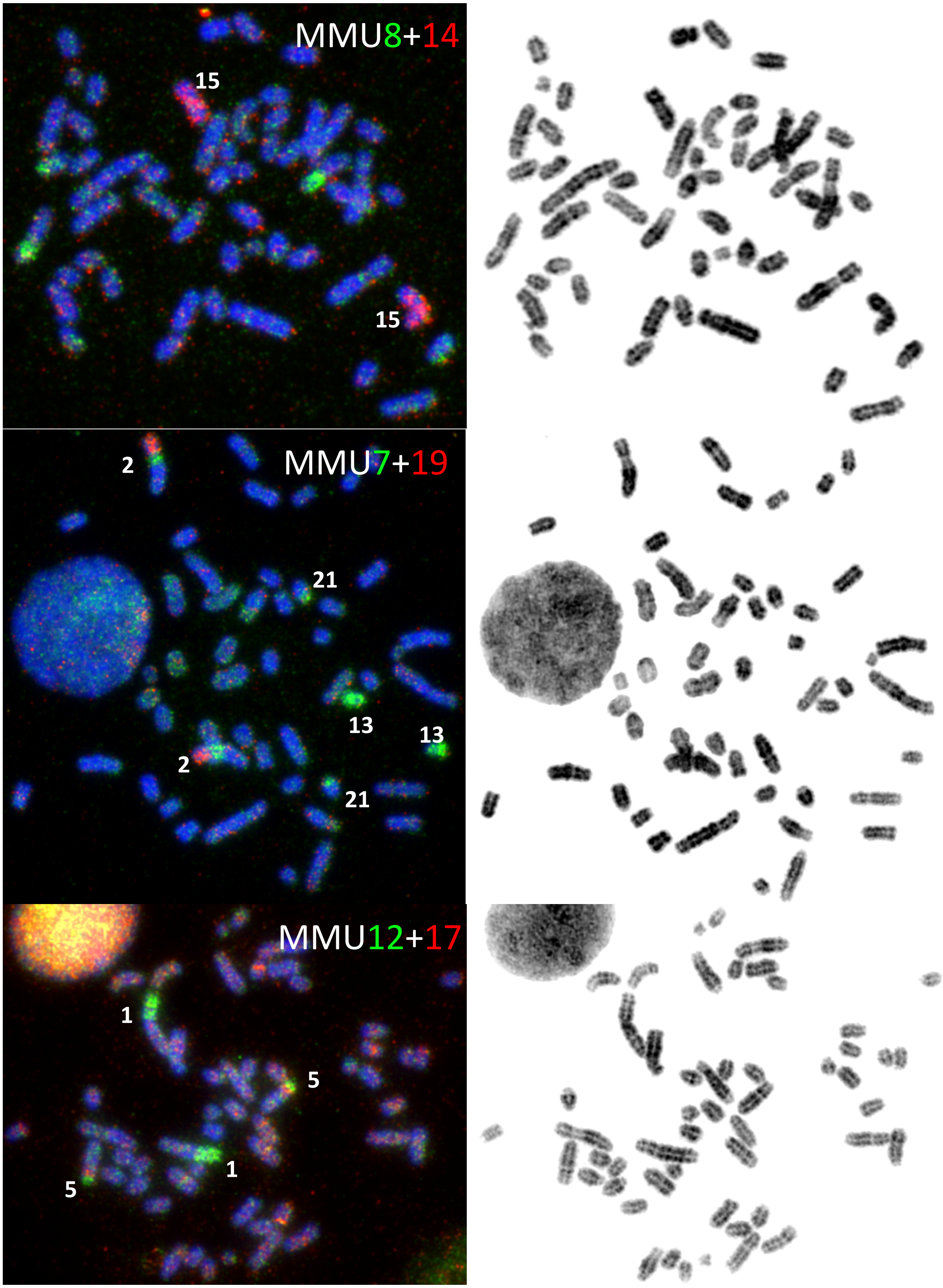
MMU8+14
15
15
MMU7+19
2
21
13
13
2
21
MMU12+17
1
5
1
5

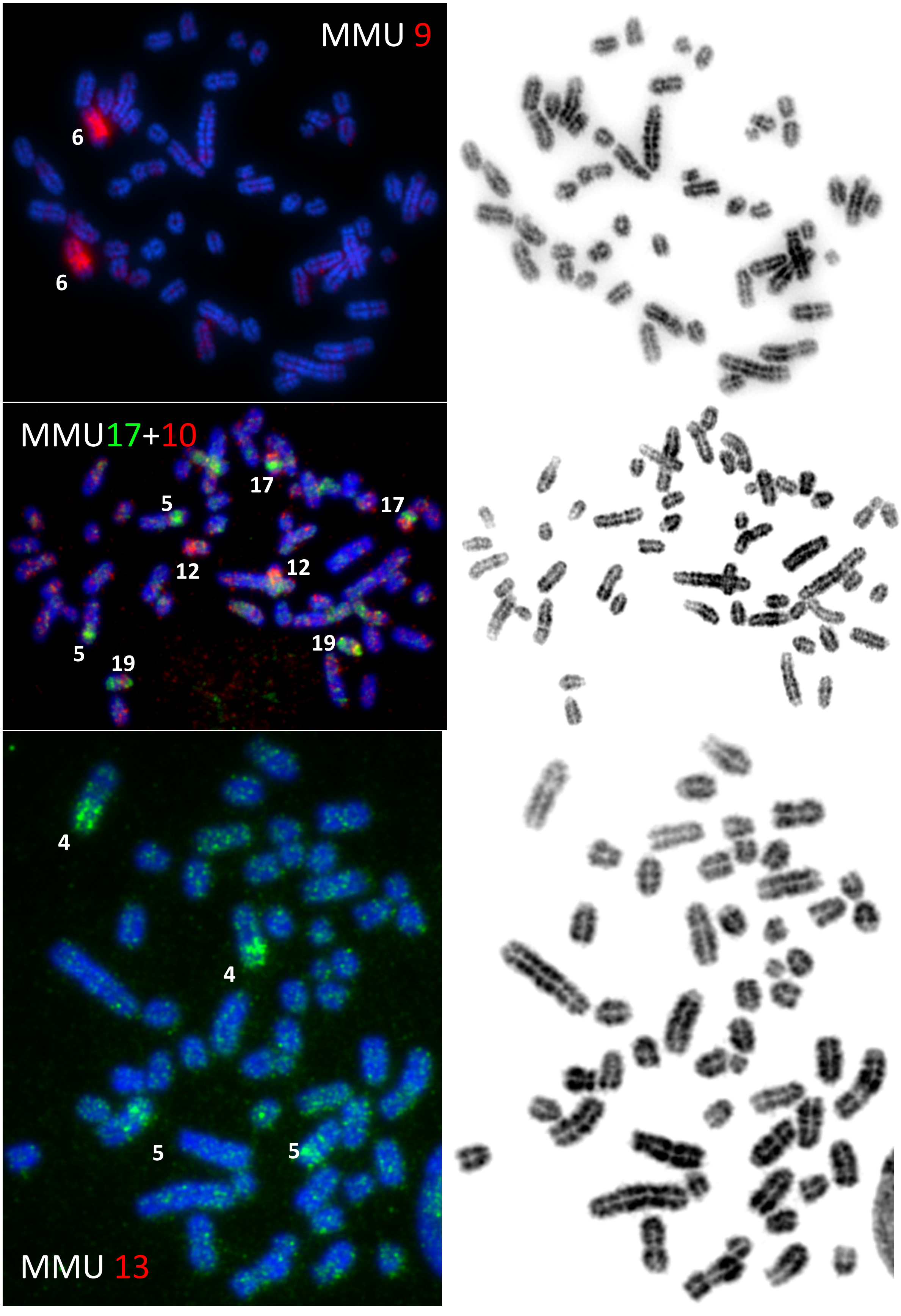
MMU 9
6
6
MMU17+10
5
17
17
12
12
5
19
19
4
4
5
5
MMU 13

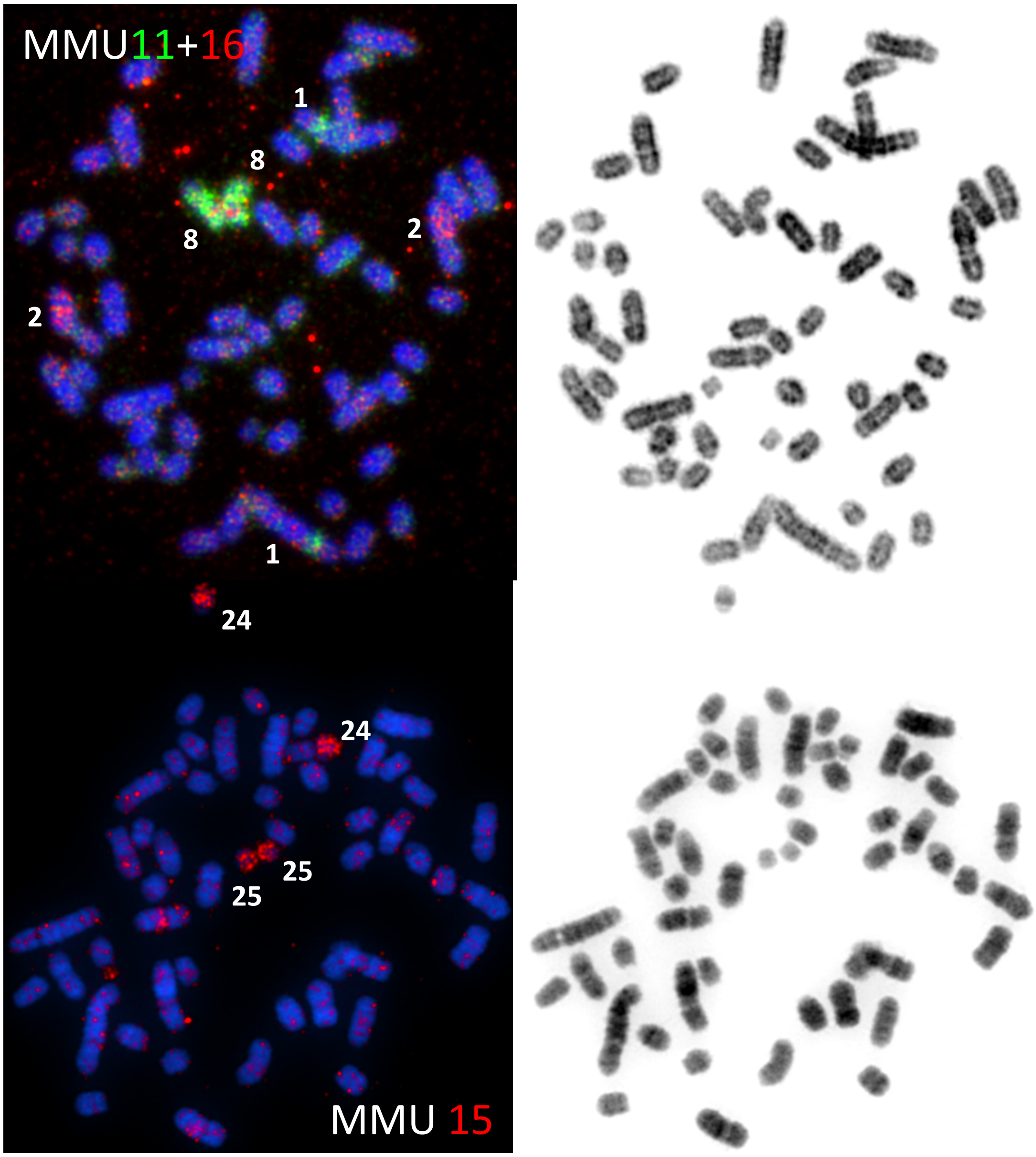
MMU11+16
1
8
8
2
2
1
24
24
25
25
MMU 15

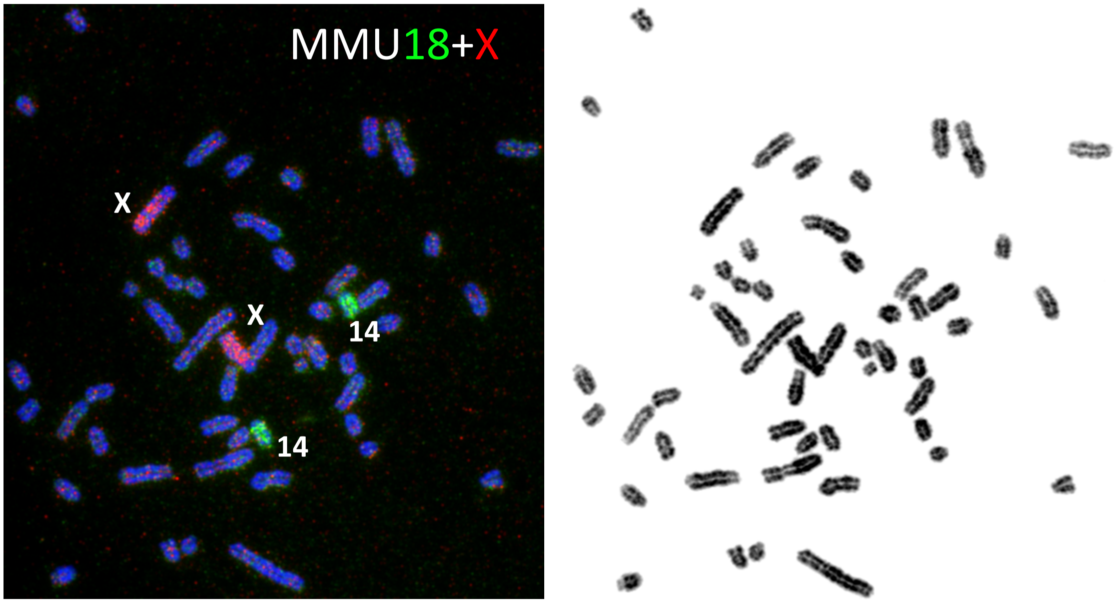

Supplementary Figure S1. **Homologous chromosomes in cotton rat and mouse.**

(a) **Mouse chromosome paints identify homologous *S. hispidus* chromosomes.** *S. hispidus* metaphase chromosomes hybridized with mouse (*M. musculus*) probes (labeled as MMU Chr. Number, *green and/or red fonts*). *Left*, fluorescent signal of probes hybridized to *S. hispidus* chromosomes (*numbered, white font*). For some spreads, multiple fluorescent channels are superimposed to represent distinct chromosomal probes marked with distinct fluorophores (*pseudocolors*, e.g. green, pink). *Right*, DAPI staining for matched karyotype to identify *S. hispidus* chromosomes.

(b)

Chromosome Homology Map Between MusMus & SigHis (*Sigmodon hispidus,* 2n = 52,XY)

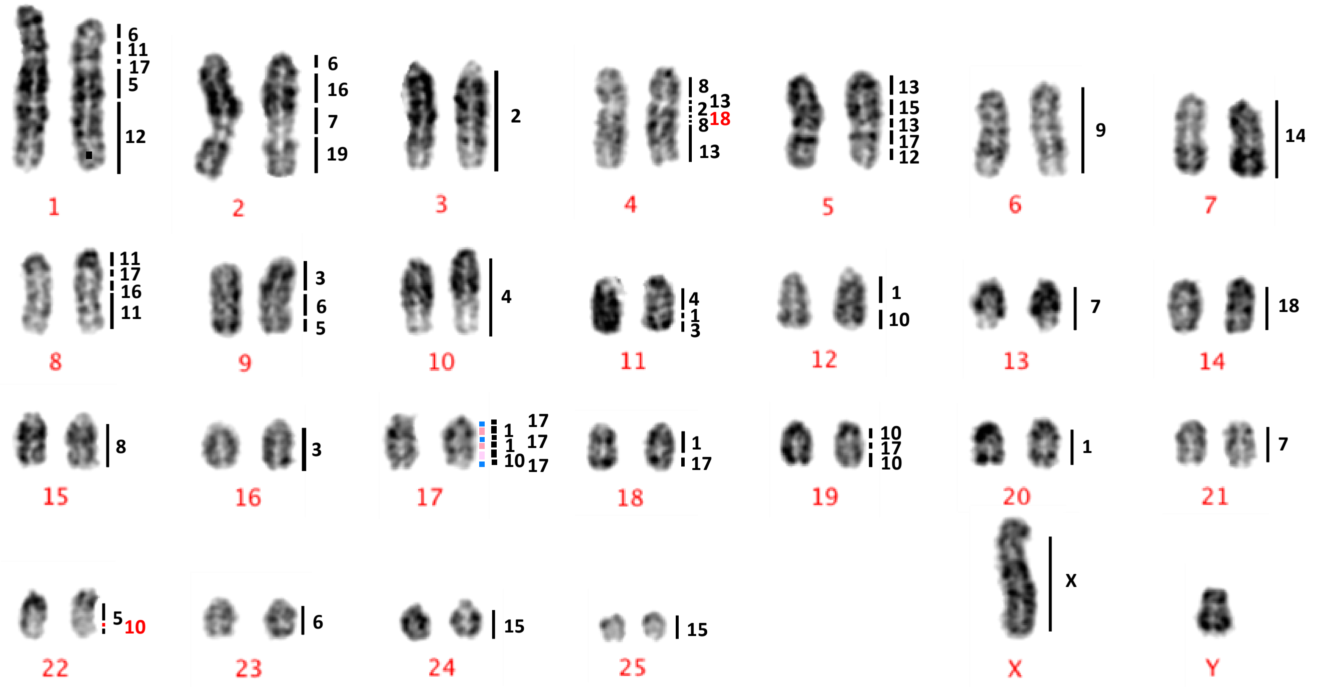

Supp. Fig. S1(b). **Chromosome homology map.** Cross-species hybridization was performed in *S. hispidus* chromosomes (*red font*) based on mouse chromosome paints (*vertical bar*, *right*). *Black font*, mouse chromosome numbers; *red font*, cotton rat chromosome numbers.

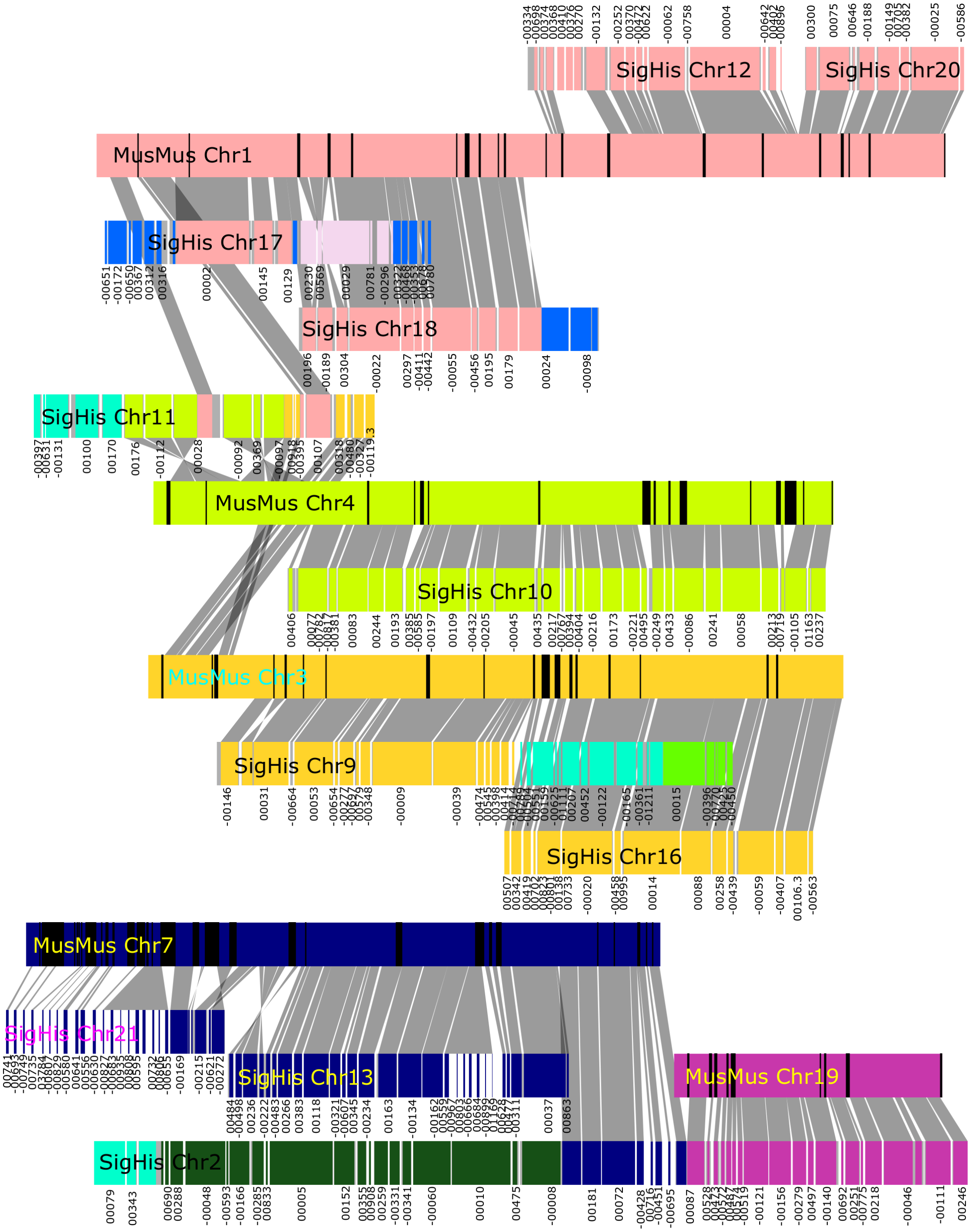
SigHis Chr12
SigHis Chr20
MusMus Chr1
SigHis Chr17
SigHis Chr18
SigHis Chr11
MusMus Chr4
SigHis Chr10
MusMus Chr3
SigHis Chr9
SigHis Chr16
MusMus Chr7
SigHis Chr21
SigHis Chr13
MusMus Chr19
SigHis Chr2

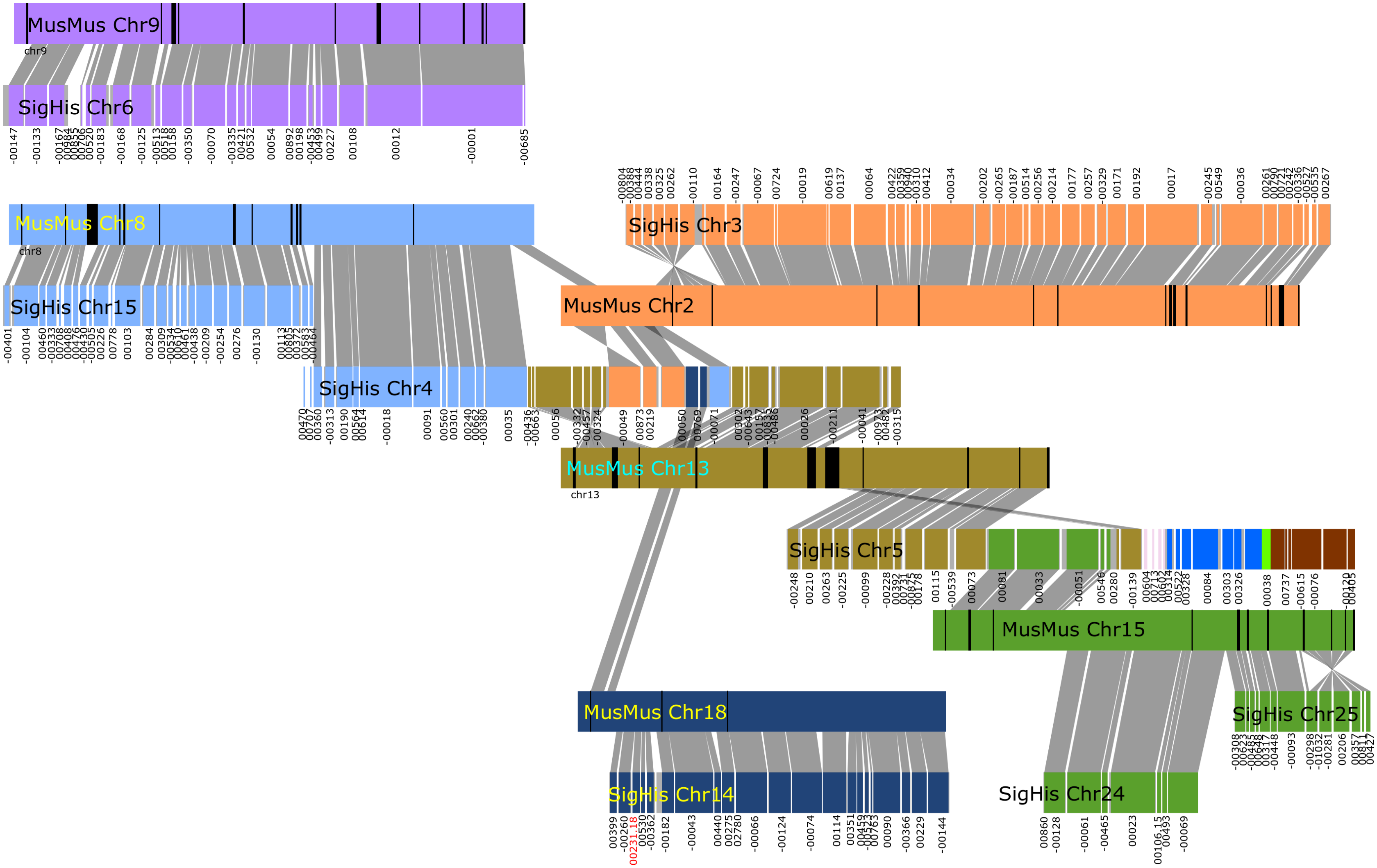
MusMus Chr9
chr9
SigHis Chr6
MusMus Chr8
chr8
SigHis Chr15
SigHis Chr3
MusMus Chr2
SigHis Chr4
MusMus Chr13
chr13
SigHis Chr5
MusMus Chr15
MusMus Chr18
SigHis Chr25
SigHis Chr14
SigHis Chr24

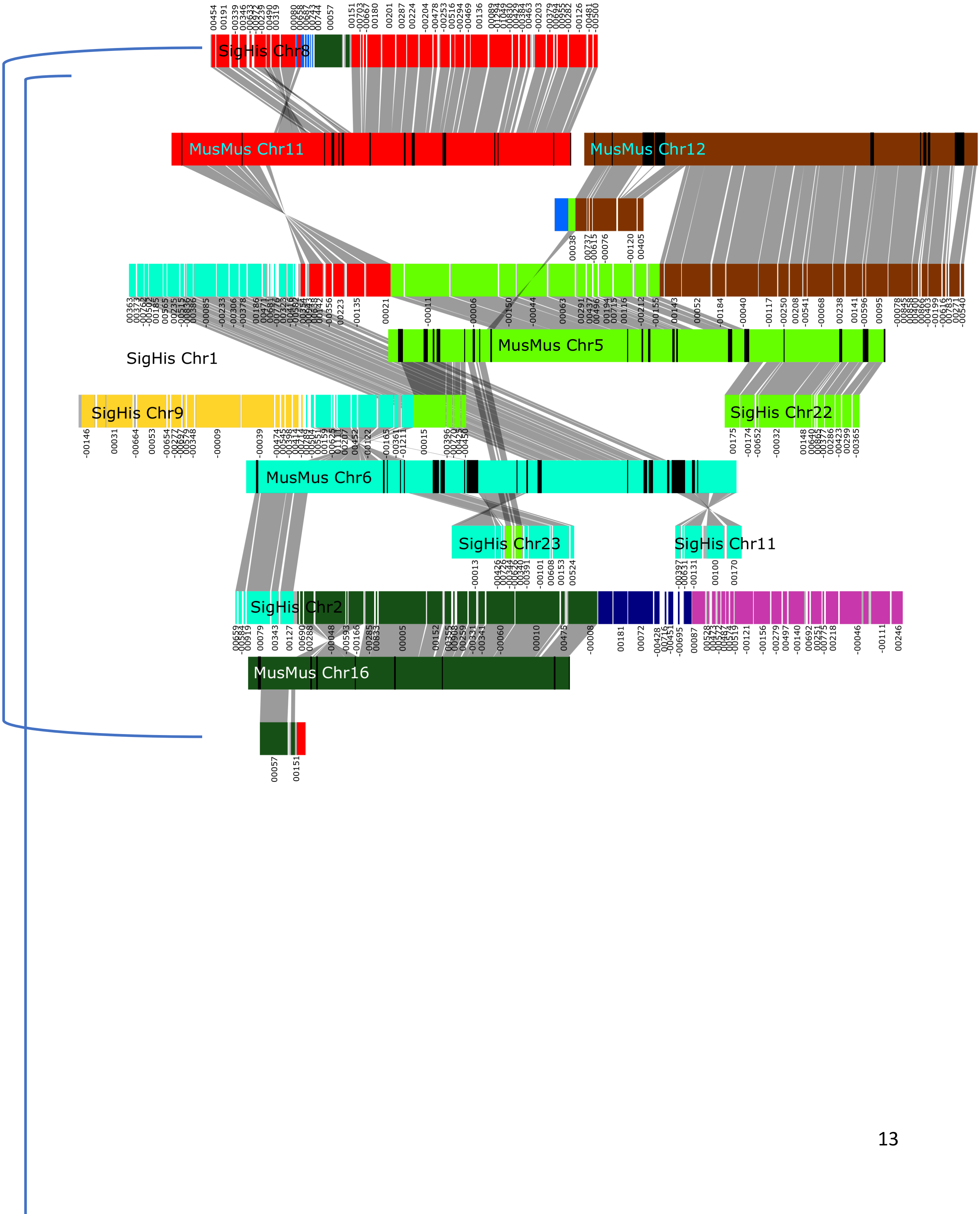
SigHis Chr8
MusMus Chr11
MusMus Chr12
SigHis Chr1
MusMus Chr5
SigHis Chr9
SigHis Chr22
MusMus Chr6
SigHis Chr23
SigHis Chr11
SigHis Chr2
MusMus Chr16

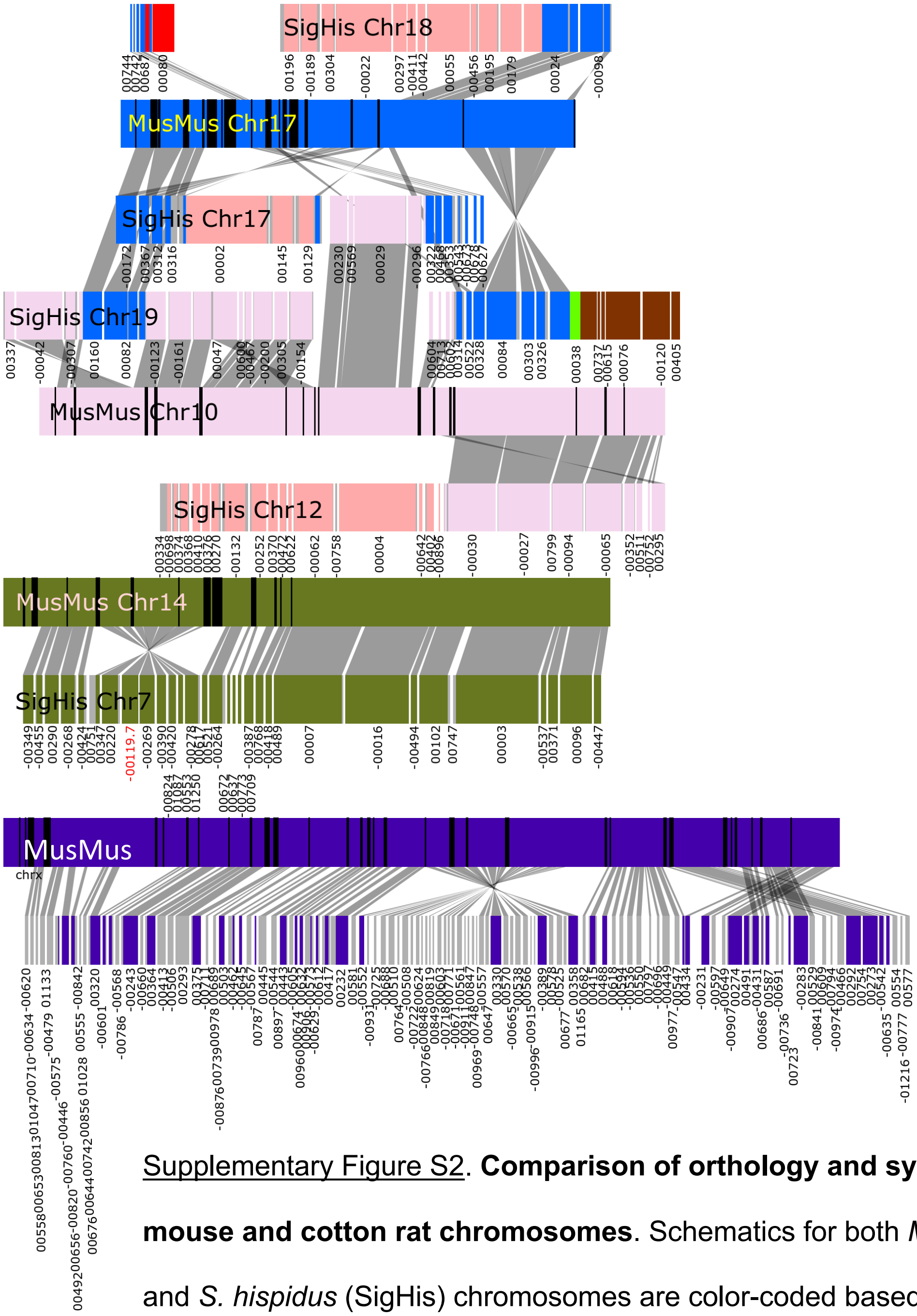

Supplementary Figure S2. **Comparison of orthology and synteny between mouse and cotton rat chromosomes**. Schematics for both *M. musculus* (MusMus) and *S. hispidus* (SigHis) chromosomes are color-coded based on *Mus musculus* chromosome assignments, as per **Figure 1**. *Gray connectors*, aligned segments display orthology and synteny. *Numbers under SigHis chromosomes*, scaffold names from de novo assembly. *Minus (-)*, reverse complemented scaffolds. *Red numbers*, disjoint scaffolds broken during pseudo-chromosome assembly. *Black stripes on MusMus chromosomes*, strain-specific diverse regions (SSDRs) containing a high density of repeats (Lilue et.al 2018).

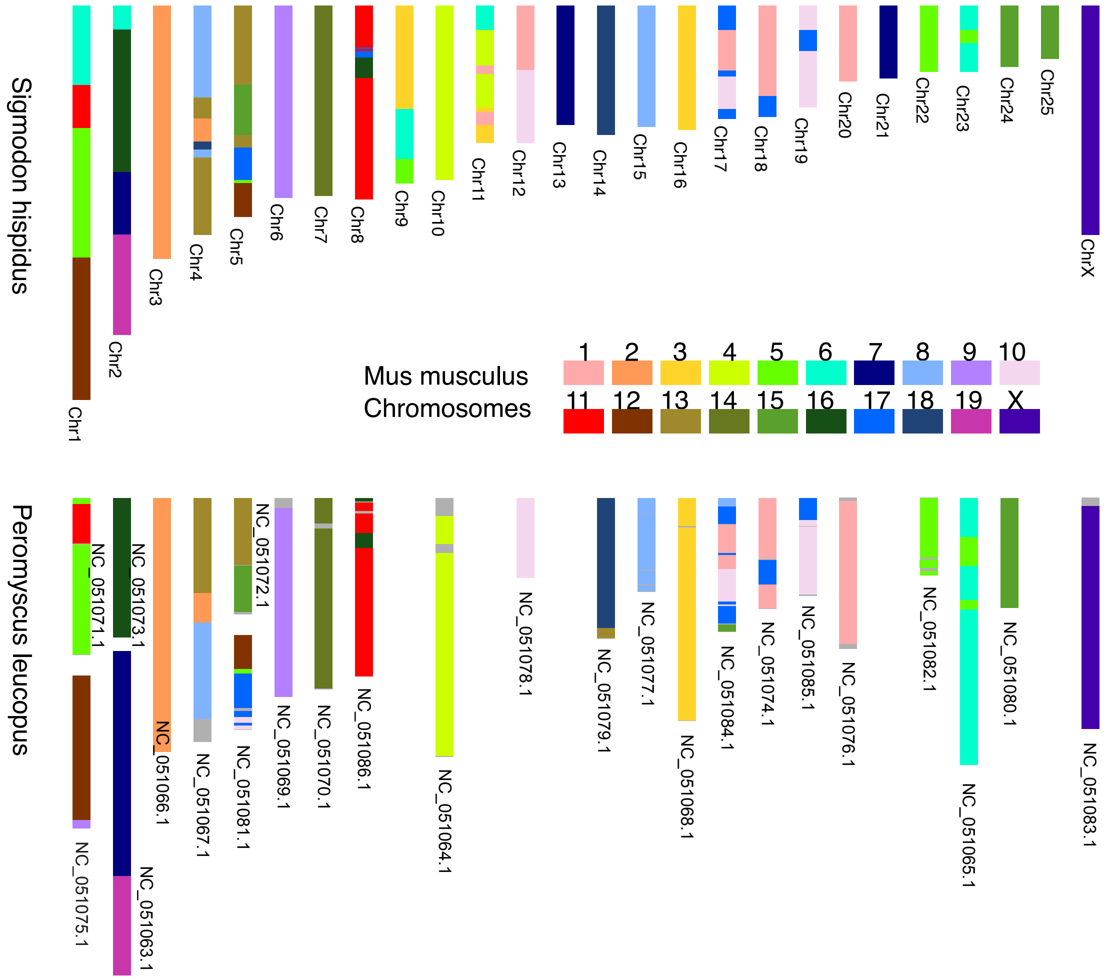

Supplementary Figure S3. **Comparison of pseudo-chromosome assemblies of cotton rat (*S. hispidus*) and white-footed mouse (*P. leucopus*) reveals shared orthology with mouse.** Schematics represent karyotypes from metaphase spreads of (*top*) cotton rat (*S. hispidus*) and (*bottom*) white-footed mouse (*P. leucopus*), painted with probes derived from *M. musculus* chromosomes (*key, colors*). Color codes on *S. hispidus* chromosomes are based on *Mus musculus. Bottom, P. leucopus* chromosome sequences were annotated previously; National Center for Biotechnology Information (NCBI) accession numbers are assigned as indicated.

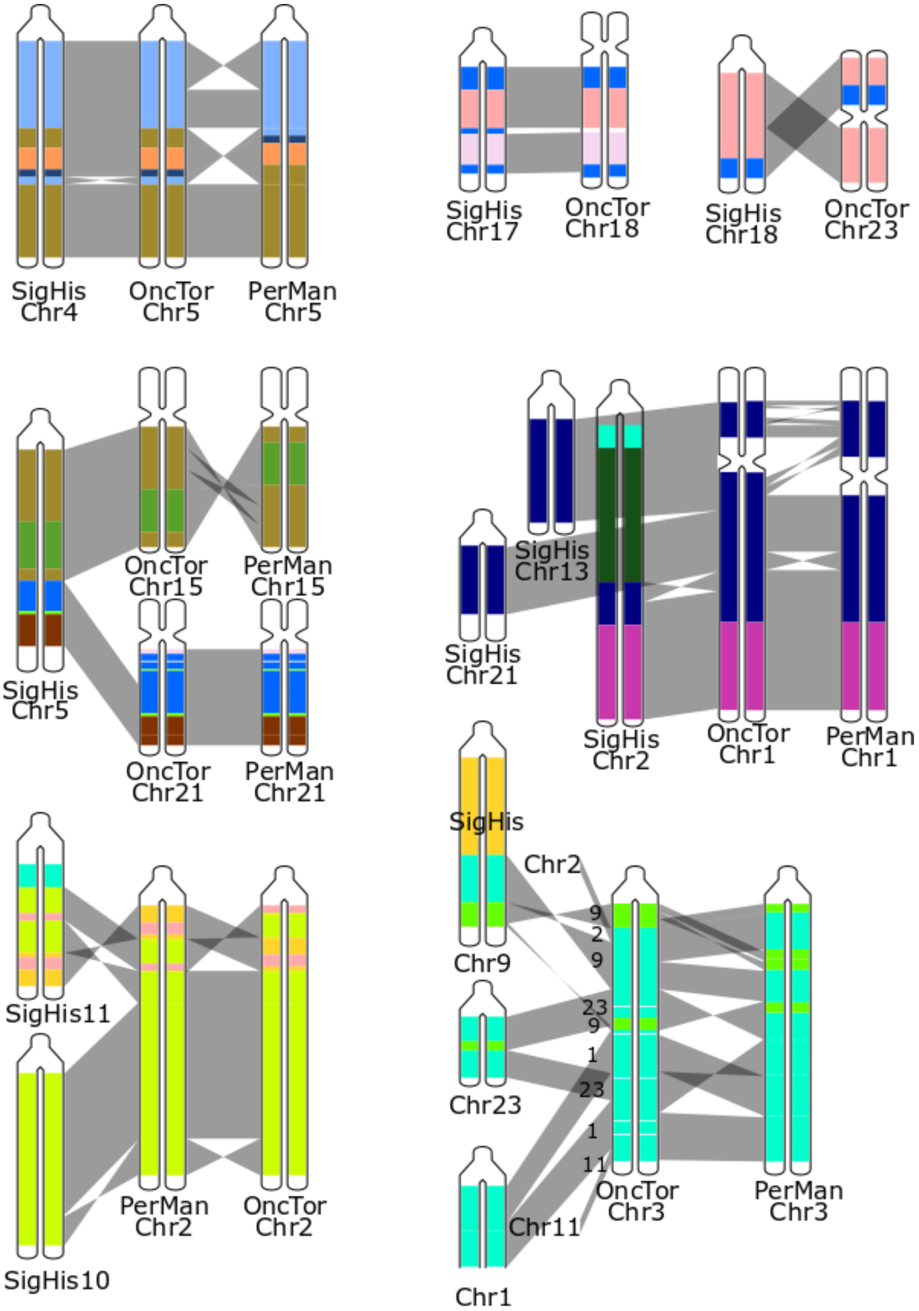

Supplementary Figure S4. **Comparison of orthology and synteny between *S. hispidus, O. torridus* and *P. maniculatus* chromosomes.** Shown are schematics for *S. hispidus* (SigHis) Chromosomes 2,4,5,9,10,11,13,17,18, 21 and 23, with syntenic alignments indicated for *O. torridus* (OncTor) and *P. maniculatus* (PerMan). Compared with mouse (*M. musculus*), these New World rodent species have higher similarity to *S. hispidus*. *Colors*, based on *Mus musculus* chromosome assignments (cf. **Figure 1**).

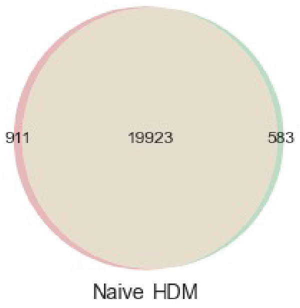

Supp. Figure S5. **Gene counts in cotton rat tissue pools.**

We pooled multiple *S. hispidus* tissues from two adult male individuals: one was exposed to house dust mite (HDM) by intraperitoneal (ip) injection followed by intranasal exposure, while the other individual was not exposed to HDM (i.e. naïve). The two tissue pools were analyzed by RNA-seq to provide a snapshot of the overall cotton rat transcriptome. Shown here is a Venn diagram enumerating genes identified by at least one RNA-seq read in at least one of the two pools, resulting in a total of 21,417 genes expressed in at least one of the pools, and 19,923 genes detected in both.

### ALR family

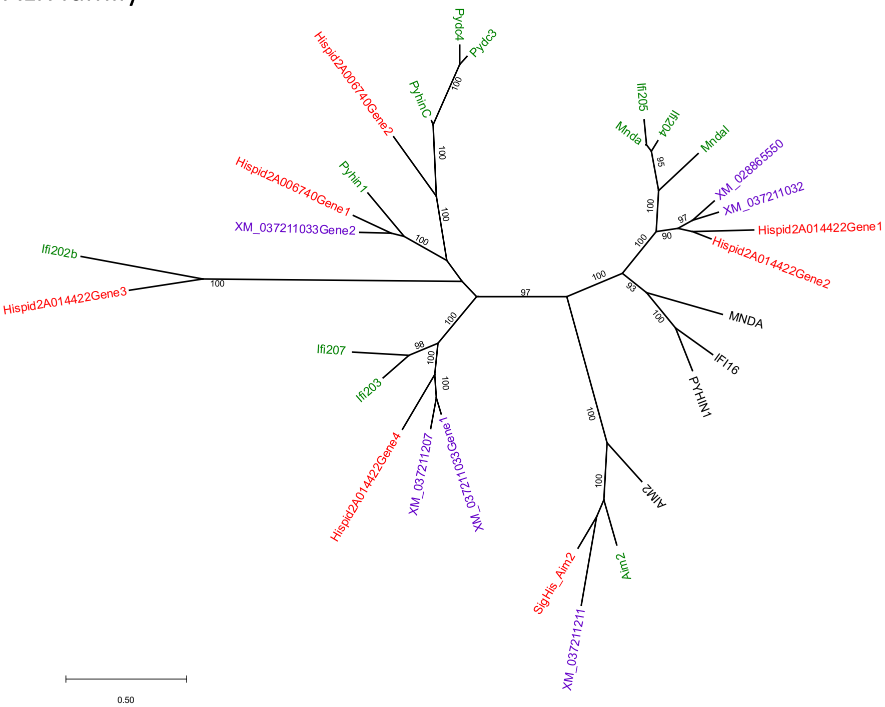

### ANG family

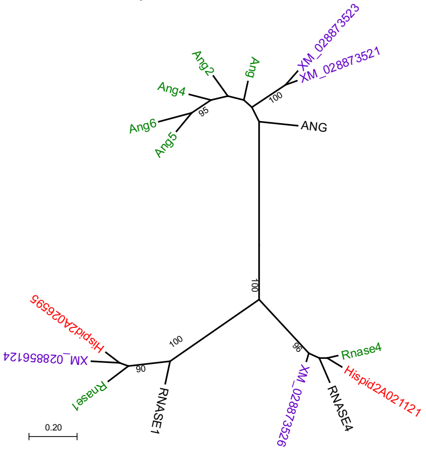

### APOL family

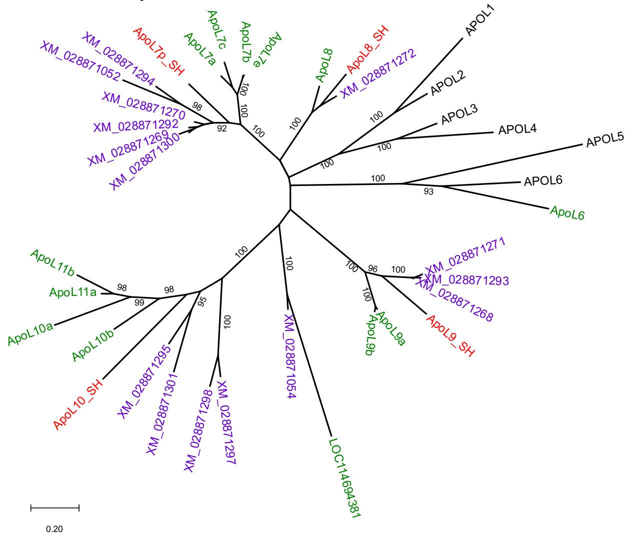

### CCL family

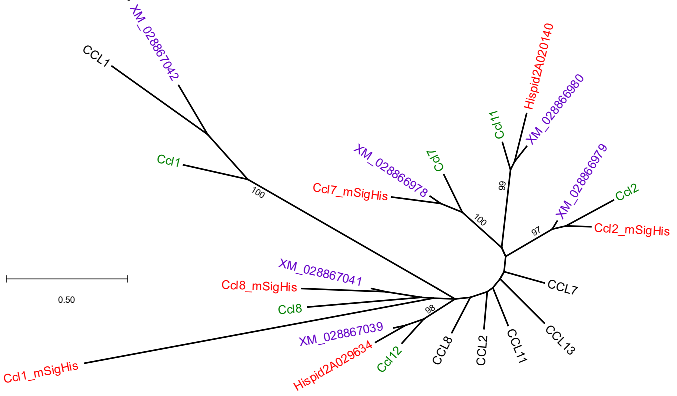

### CD300 family

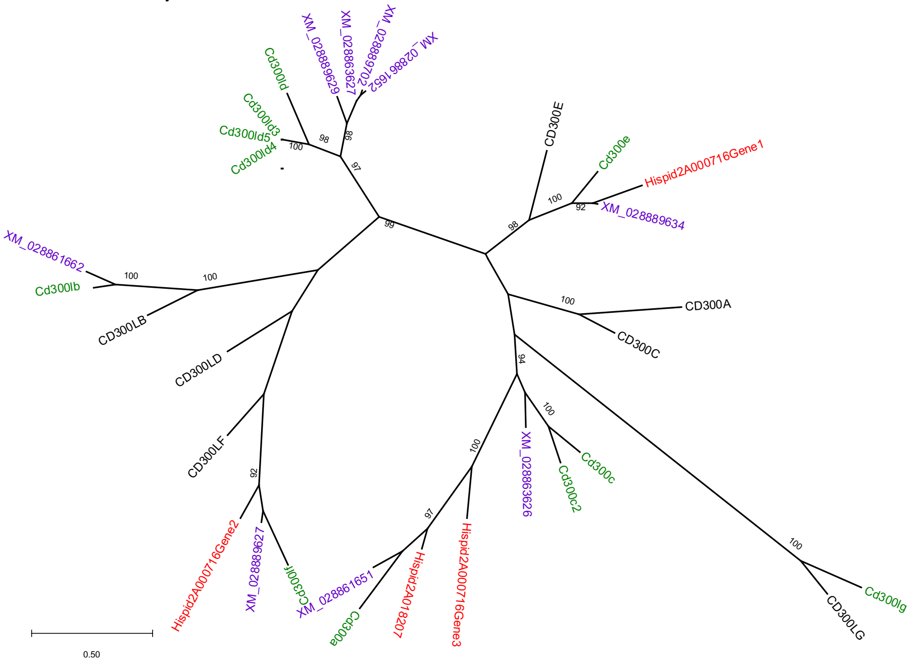

### CXCL family

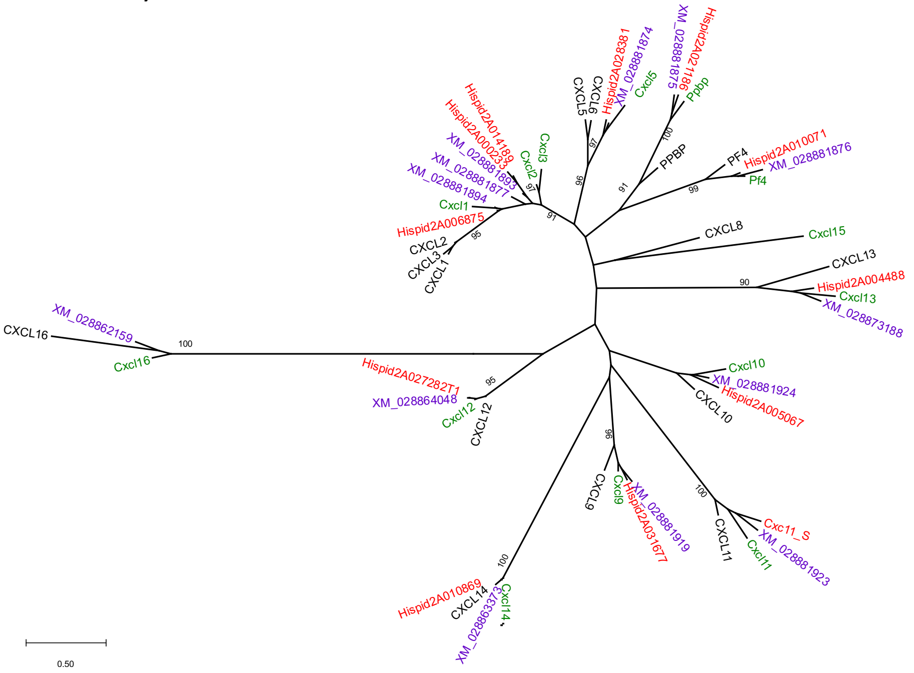

### CXCR family

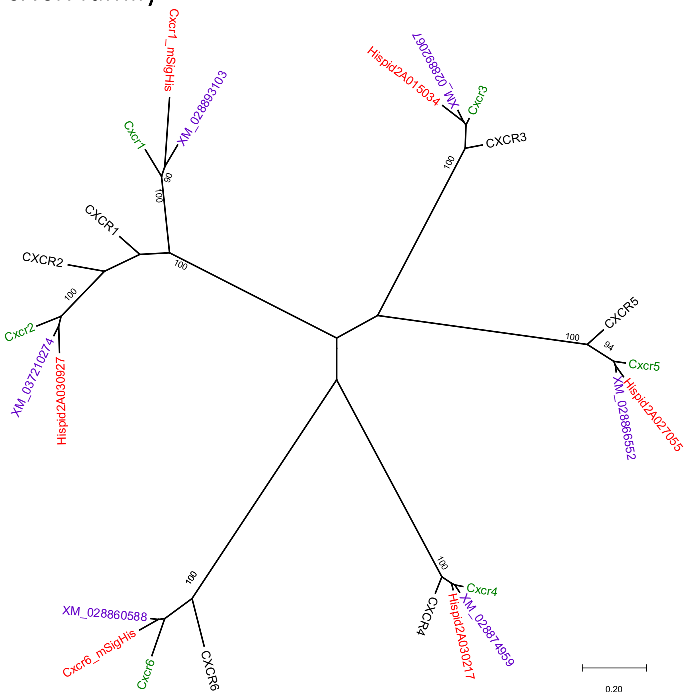

### GBP family

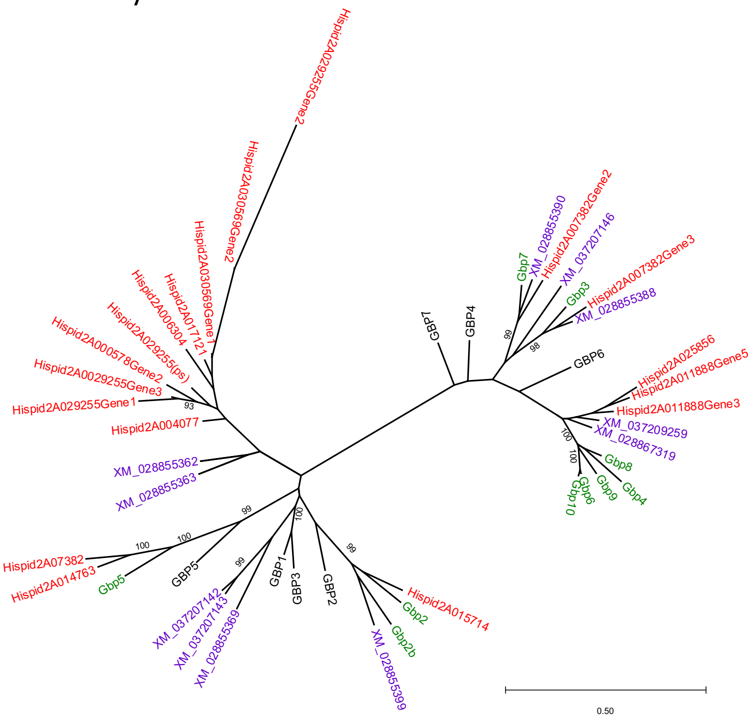

### GSDM family

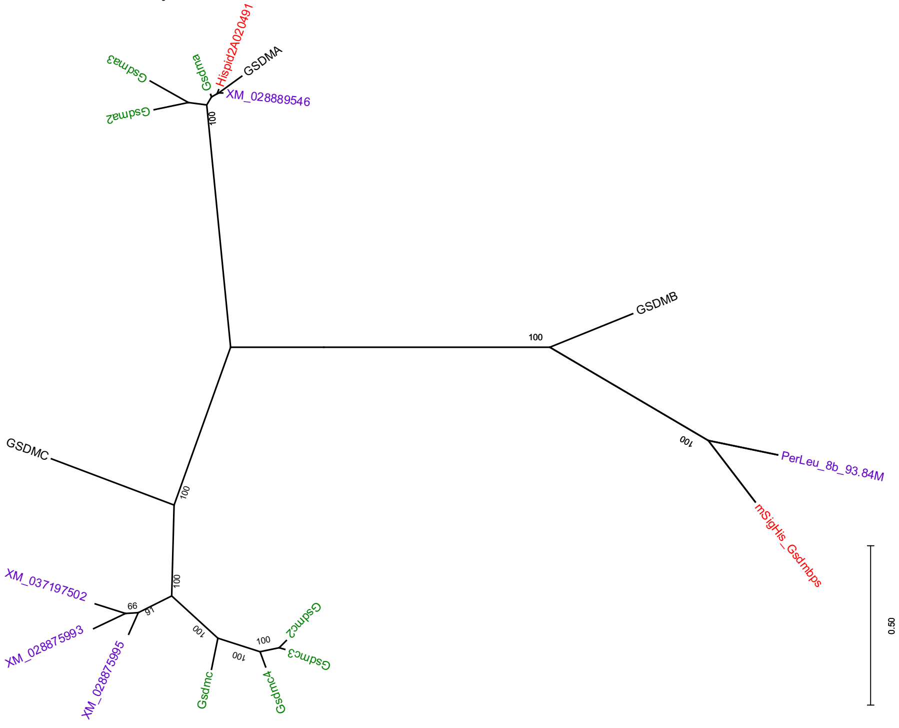

### Ifit family

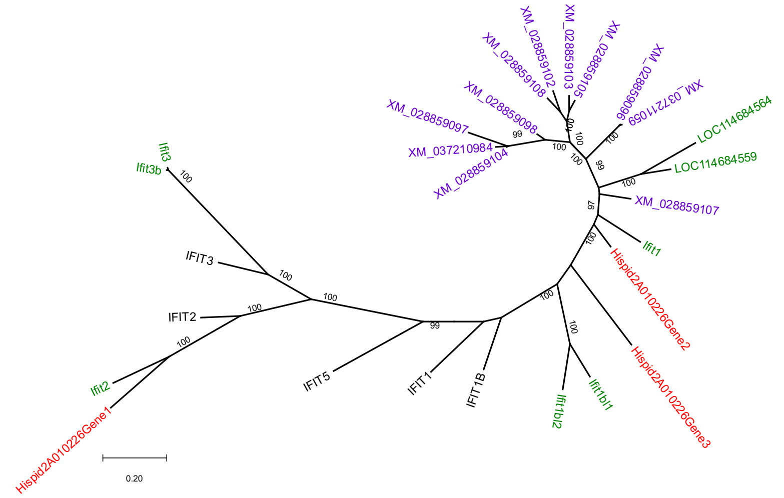

### IFITM family

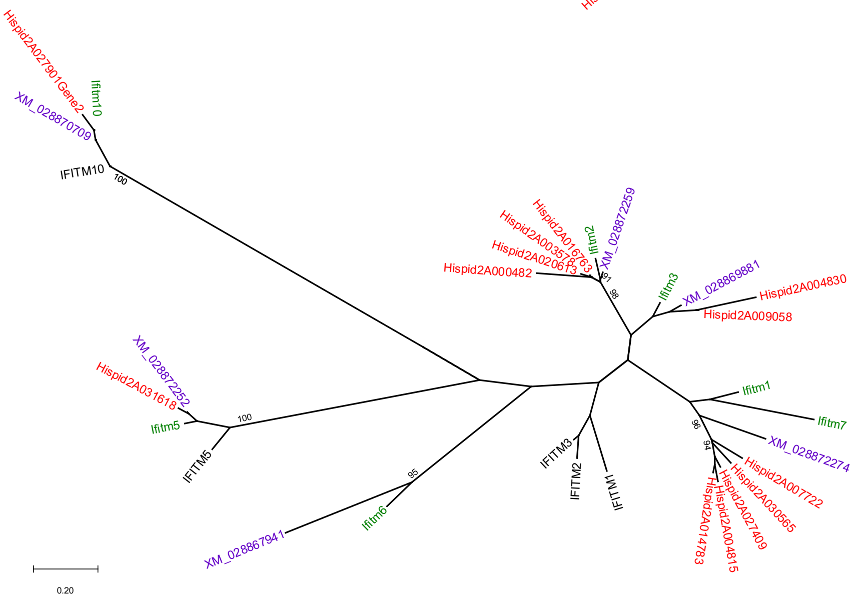

### IRG family

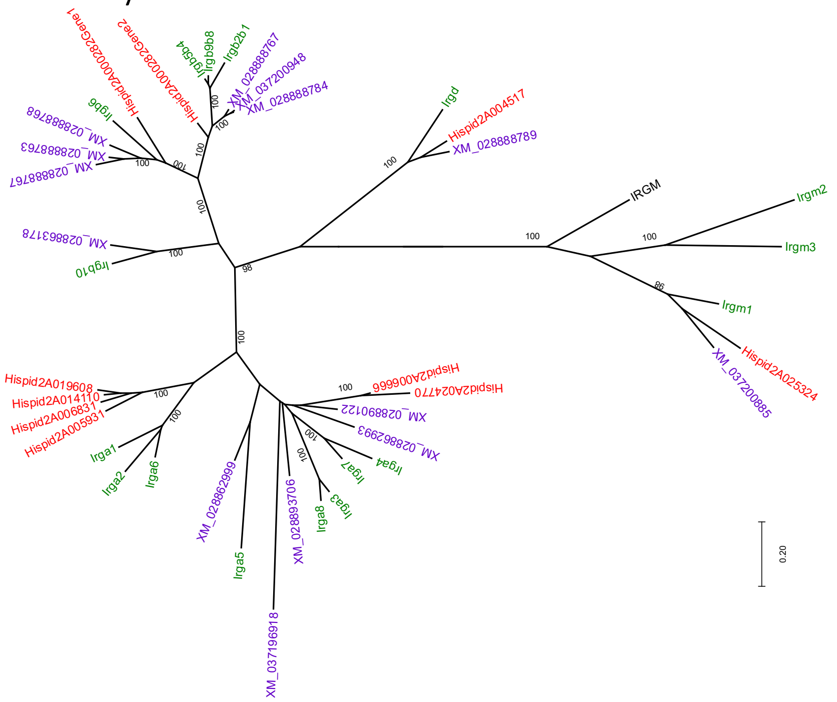

### MX family

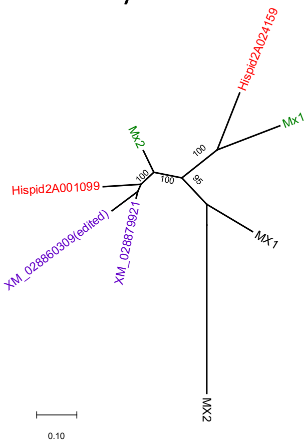

Nlrp family

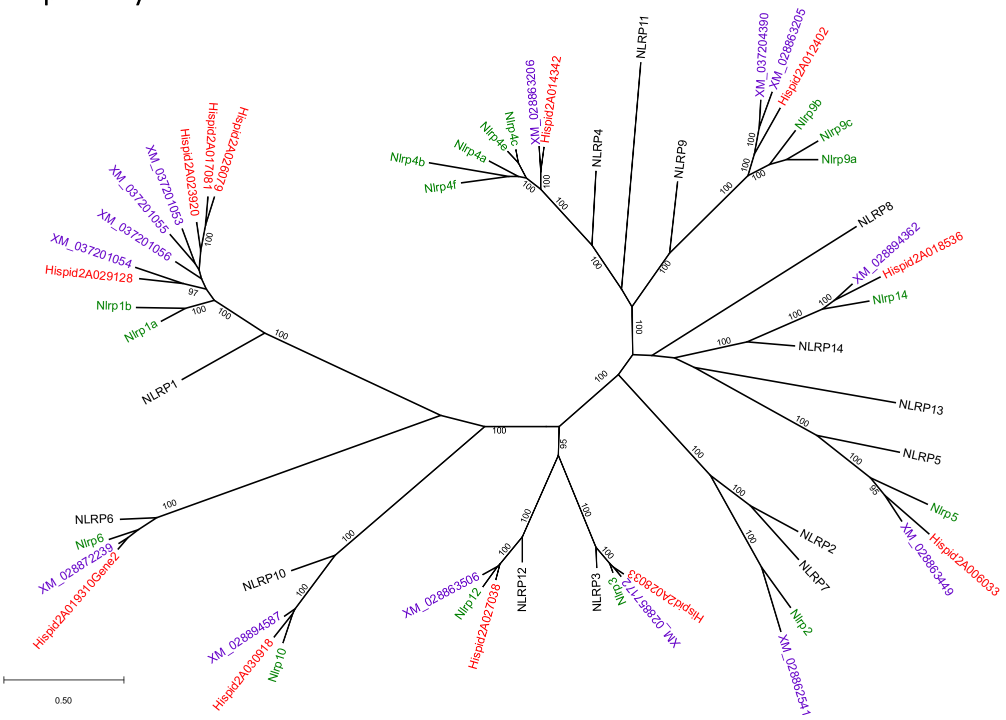

Oas family

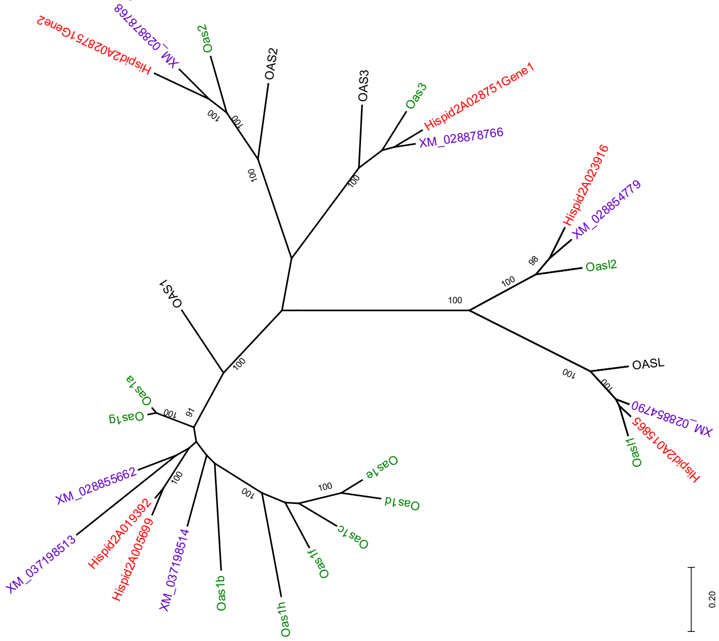

### Raet / Ulbp family

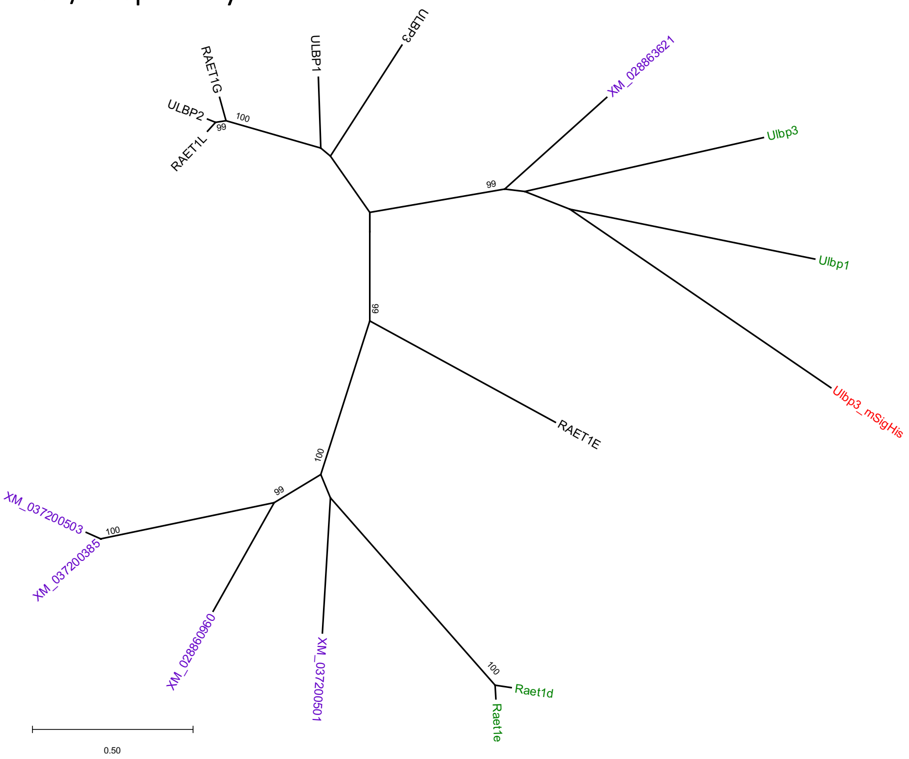

### Slfn family

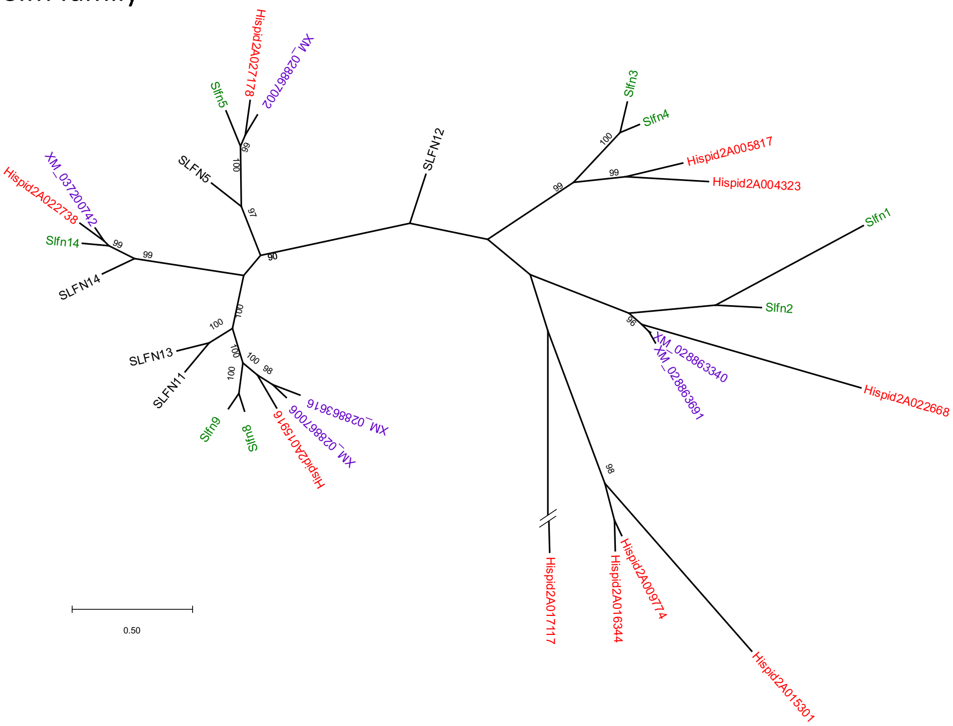

Supplementary Figure S6. **Phylogenetic trees of 17 immune-related gene families.** Coding sequences of each gene were aligned. Nucleotide maximum likelihood trees were built with Mega X (Kumar et al. 2018). In each tree, green, purple, red and black branch indicate *M. musculus* (GRCm38 reference), *Peromyscus leucopus*, *S. hispidus* and human genes, respectively. Bootstraps are shown when > 90.

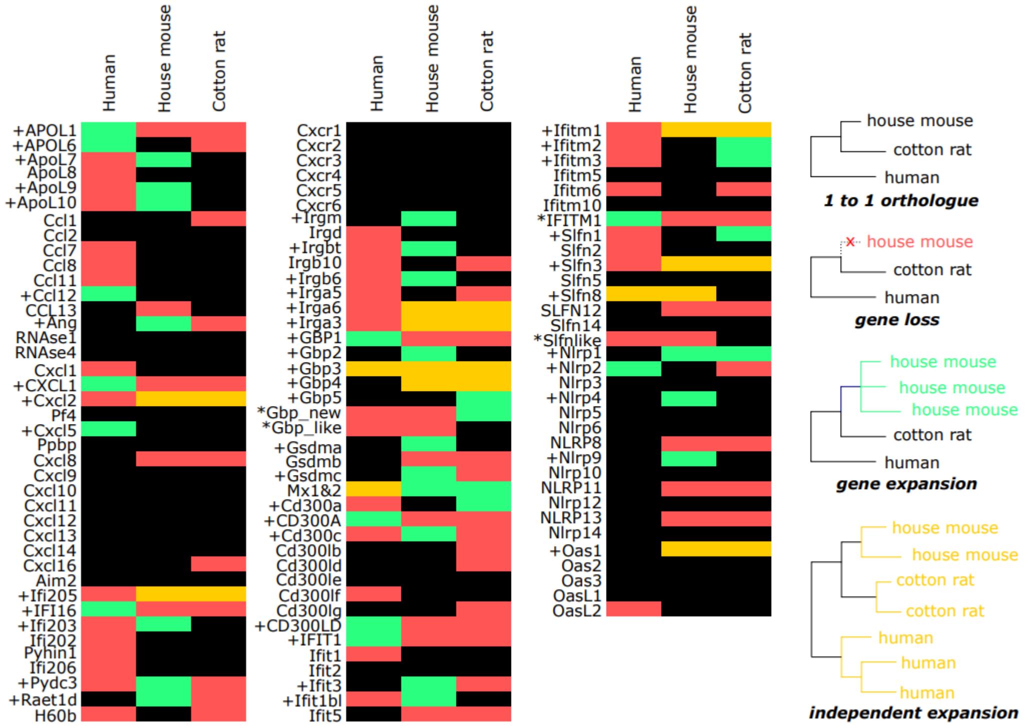

Supplementary Figure S7. **Three-way comparison of 352 immune-related genes from 17 families.** *Black*, *red*, *green* and *yellow*: direct orthologs, gene loss, gene duplication and species-specific gene expansion, respectively; “+” in front of gene names, multiple mouse or human genes included in gene duplication; and “*”, comment on the gene. See Supplementary Table S2 for more details.

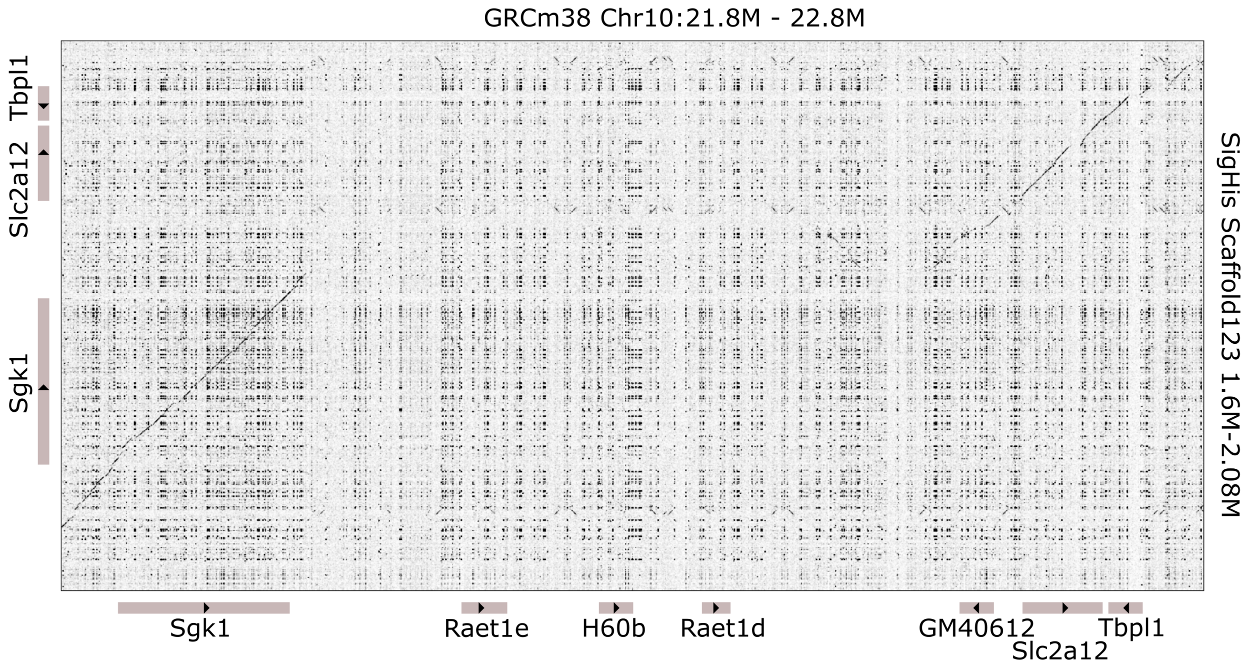

Supplementary Figure S8. **Loss of *Raet* / *MHA* locus from cotton rat.** This dot plot compares homology or identity at the mouse Chr10: 21.8M - 22.8M genomic locus (*x-axis*) against our *S. hispidus* de novo assembly (*y-axis*). *Black dots*, homology between the two species' genomes; *diagonal lines*, aligned, contiguous genomic sequences showing homology. A deletion of ~ 500 kb (*gap in diagonal line*) of the mouse genome (*x-axis*, mouse reference genome; *bottom*, annotated genes) is observed in (*y-axis*, Sig His, *left*, annotated genes) cotton rat, between genes *Sgk1* and *Slc2a12*, causing losses of *Raet1e*, *H60b*, *Raet1d* and possibly *GM40612*.

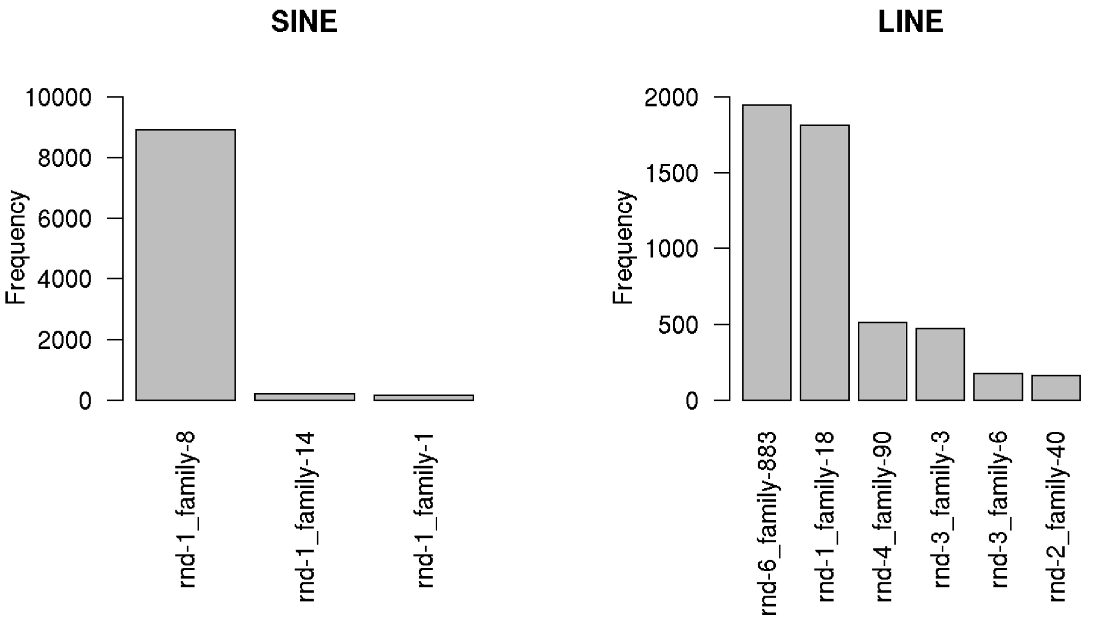

<u>Supplementary Figure S9.</u> **Counts of repetitive elements by family.**

We found 16,384 repetitive elements with less than 2% sequence diverence from consensus sequences. Most of these young elements are SINE or LINE elements. We counted the number of elements by repeat family for SINE and LINE classes. (Left) Number of young elements by SINE family. We show three SINE families having 100 or more elements with <2% diversity from the consensus in cotton rat genome. One family (rnd-1-family-8#SINE/B2) represents most of the young SINE elements (8910 out of 9676 elements, 92%). (Right) Number of young elements by LINE/L1 family. We show six L1/LINE families having 100 or more elements with <2% divergence from the cotton rat genome consensus. Although RepeatModeler called these six families as independent families, five families (rnd-1_family-18, rnd-4_family-90, rnd-3_family-3, rnd-2_family-40, rnd-3_family-6) are 5' truncated sequences of the sixth (rnd-6_family-883#LINE/L1), which therefore we designate as the consensus cotton rat L1 family.

```
>rnd-1_family-8#SINE/B2
GGGGCTGGAGAGATGGCTCAGCGGTTAAGAGCACCGTCTGCTCTTCCAGAGGACCCGGGT
TCGATTCCCAGCACCCACATGGCAGCTAACAACTGTCTGTAGTTCCAGTTCCAGGCAAAC
ACCAAGGCAAATAAATAAATAAAGTTTAAAAAAAAAAAAAAAAAAAAAAAAAAAAAA
```

Supplementary Figure S10. **Nucleotide sequence of consensus cotton rat SINE B2 element**.

This rnd-1_family-8#SINE/B2 = SINE B2 family is the most frequently detected young SINE family in the cotton rat genome. This family is closest to the mouse SINE B3 family at 9.6% divergence.

```
>CottonRat_L1_ORF1 (486 nt|161 aa)
MFINKIIEENFPNLKKEMPIKIQEAYRTPNRYDPXKKSARHIIIKPQNIQ
NKEKILRAAKEKGQLTYKGRPIRITPDYSMETMKARRAWSEVIQTXKQHG
CQTRILYPAKLSITIDGVNKVFQDKVRLKEYLSTNPALQRVLEGKLQPKD
PRYTHEITNNQ
>CottonRat_L1_ORF2 (3825 nt|1274 aa)
MAGNMNQWSLISLNINGLNSPIKRHTLTRWIQQQKPSFCCLQETHLNCKD
RHRLRIKGWEKIFQSNGPKKQAGVAILISNKVDFKLKAXKRDQEGHYLFI
TGKIGQEEISILNIYAPNIGAPTYVKETLLKLESHINSYTLIVGDFNTPL
STLDRSTRQSLNREIRSLNEVMMQMDXIDIYRTFHPKAKEYTYFSAPHGT
FSKIDHIIGHKANISRYKKVEIIPCNISDHQGLKLEFNNIKTQRKPTNSW
KLNIDQLNHLWVKEEIKKEIKVFLEFNENESTTYPNLWDTMKAVLRGKFI
ALDAYMKNLEKSHTRDLTVQLKALEQLEANSPRRSRRQEIIKLRAEINTI
ETKRTIQRINESKSWFFEKVNKIDKPLAKLIKKQRESVQINKIRNEKGDI
TTDPQEIQRIIRSYFENLYSTKMENLEEMDSFLDRHHLPKLNEDQINYLN
RPITLKEIEAVIKXLPKNKSPGPDGFSAEFYQTFKEELLPLLLKLFHTVE
TEGILPNSFYEATITLIPKPLKDTTKKENYRPISLMNIDAKILNKILANR
IQQCIKDIIHHDQVGFIPEMQGWFNIRKTVNVIHHXNKLKEQNHMIISLD
AEKAFDKIQHSFMIKALERSGIKGEYLNIIKAIYKKPTASIKLNGEKLKE
IPLKSGTRQGCPLSPYLFNIVLEVLAKAIRERKEIKGIQIGKEEVKLSLF
ADDMIVYISDPQNSSKELLQLINTFSNVAGYKINSKKSVALLYAKDRRAE
KEIRETLPFILAANSIKYLGVTITKEVTDLFDKNFKSLKKEIEEDIRKWK
ALPCSWIGRINIVKMAILPKAIYRFNALPIKIPTXYFLXLERTILNFIWN
NRKARIAKTILLNKRTSGGITIPDFKLYYRATVIKTAWYWHKNRLVDQWN
RIEDPDINPKTYEHLIFDKEAQNIQWKKDSIFNKWCWHNWKEVCRKMQXD
LSLSPCTKLKSKWIKDLNLTPTTLNLIEEKVGNTLLQIGTGYHFLNITPV
AQALRPTINKWDLLKLGSFCKAKDTVNRTKRQPTEWEKIFTNPTSDRGLI
SKVYKELKNLDPKTLNNPIKKWGSDLKREFSAEELRMAERHLKKCSSSLA
IREMQIKTTLRYHLTPVRLAKIKGTDDSLCWRGCGARGXLLHCWWDCKLV
QPLWNTIWRFLRKLRINLPKDPAIPLLGIYPKEAHSHSKGISSTMFIAAL
FVIARTWKQPRCPSTEEWINKLWYIYTMEYYSAVRKNDILKFAGKWMDLE
NILLSEVSQTQRDKQGMYSLTSGL
```

Supplementary Figure S11. **Predicted amino acid sequences of consensus cotton rat L1 ORF1 and ORF2.**

As observed in every other mammalian L1 retrotransposon, the cotton rat L1 consensus sequence encodes two open read frames (ORFs).

(*Top*) Cotton rat L1 ORF1 (nucleic acid binding protein). This amino acid sequence displays 67% identity to rat L1 ORF1 (CAA37644.1), 67% identity to mouse L1 ORF1 (AAA39397.1), 70% identity to Chinese hamster L1 ORF1 (ERE72641.1). (*Bottom*) Cotton rat L1 ORF2 (encoding a reverse transcriptase). This amino acid sequence displays 74.63% identity to rat L1 ORF2 (AAY88216.1), 73.06% identity to mouse L1 ORF2 (AAC72793.1), and 62.4% identity to human L1 ORF2 (B34087).

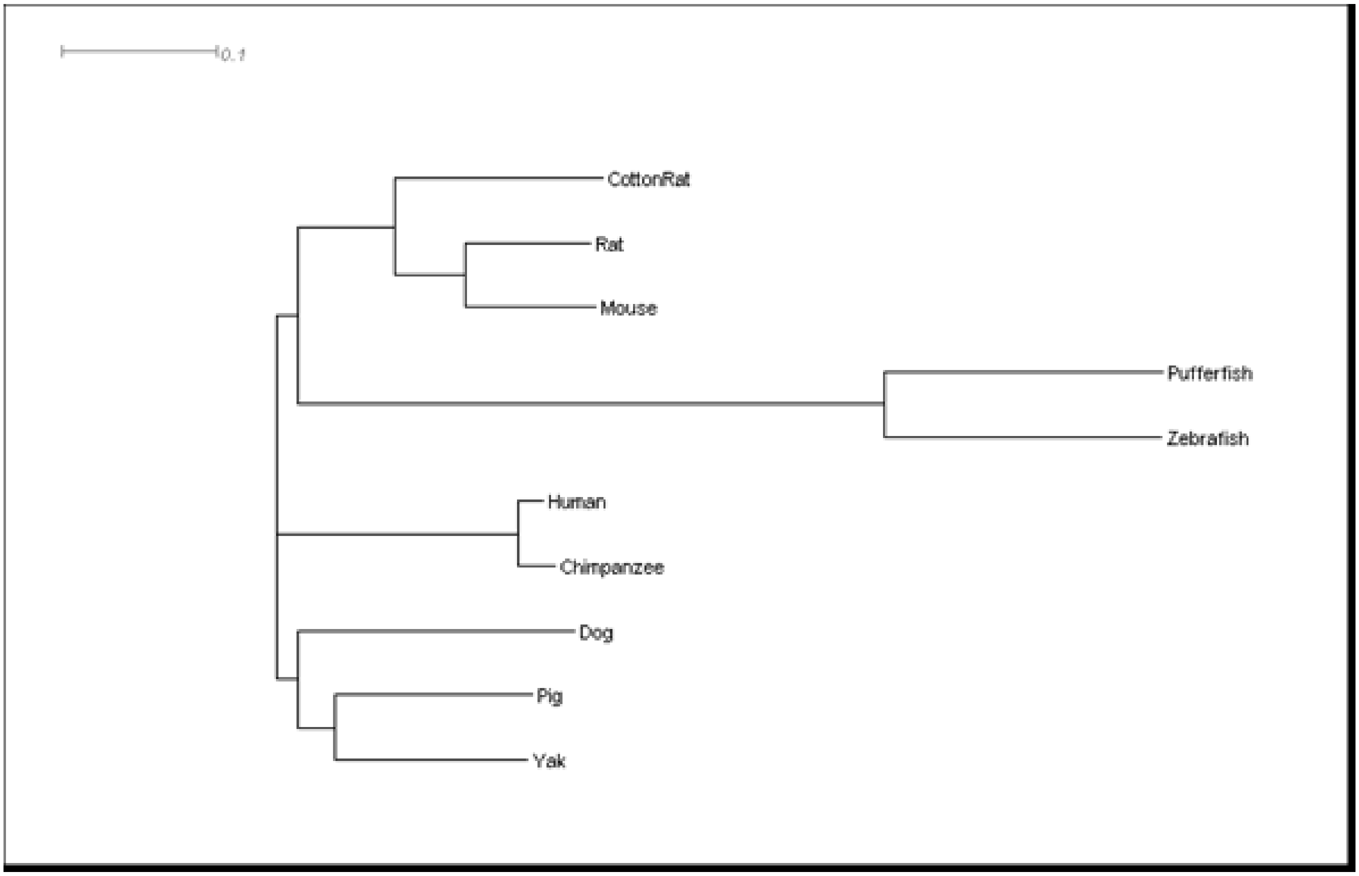

Supplementary Figure S12. **Phylogenetic tree characterizes evolution of vertebrate L1 ORF2s**

Shown here is a phylogenetic tree characterizing evolution of the L1 ORF2 (reverse transcriptase) sequences from ten vertebrate species, of which nine were obtained from NCBI protein database (Rat=AAY88216.1, Mouse=AAC72793.1, Human=O00370, Chimpanzee=ABE73458.1, Dog=BAA25253.1, Pig=ABR01162.1, Yak=ELR58410.1, Pufferfish=BAD04858.1, Zebrafish=BAC82621.1). The tenth is from our consensus cotton rat L1 sequence. Multiple sequence alignment was performed using Clustal Omega, from which a neighbor-joining tree was generated as described (Sievers et al. 2011).

**SUPPLEMENTAL TABLES**

| | |
|---|---|
| Number of input reads | 218,393,360 |
| Average input read length, nt | 283 |
| UNIQUE READS: | |
| Uniquely mapped reads, number | 186,923,111 |
| Percentage reads that are uniquely mapped | 85.59% |
| Average mapped length, nt | 281.51 |
| Number of splices: Total | 135,908,927 |
| Number of splices: Annotated (sjdb) | 0 |
| Number of splices: GT/AG | 134,915,009 |
| Number of splices: GC/AG | 702,540 |
| Number of splices: AT/AC | 65,010 |
| Number of splices: Non-canonical | 226,368 |
| Mismatch rate per base | 0.20% |
| Deletion rate per base | 0.00% |
| Deletion average length | 1.57 |
| Insertion rate per base | 0.00% |
| Insertion average length | 1.35 |
| MULTI-MAPPING READS: | |
| Number of reads mapped to multiple loci | 15,875,474 |
| % of reads mapped to multiple loci | 7.27% |
| Number of reads mapped to too many loci | 578,920 |
| % of reads mapped to too many loci | 0.27% |
| UNMAPPED READS: | |
| Number of reads unmapped: too many mismatches | 0 |
| % of reads unmapped: too many mismatches | 0.00% |
| Number of reads unmapped: too short | 14,941,431 |
| % of reads unmapped: too short | 6.84% |
| % of reads unmapped: other | 0.03% |

Supplementary Table S1. **Alignment features of bulk RNA-seq reads from cotton rat tissue pool P1**. Cotton rat RNA pool (P1) was derived from multiple tissues dissected from an individual exposed to house dust mite antigen. RNA-seq was performed (Methods) resulting in dataset 12_14_S29_L007_P0001. See **Table 3**. Sjdb, splice junctions database in STAR.

Supp. Table S2. **Selected "immune response" (GO term) genes in cotton rat genome.** Listed here are genomic features of 224 annotated genes identified using 2GO in cotton rat genome assembly that are associated with the GO term GO:0006955, "immune response". See linked Excel spreadsheet, "Supp. Table S2". To assess if annotations of our reference genome included immune response genes that were expressed in these pools, we selected 224 genes as identified in our reference assembly that were annotated by the specific gene ontology (GO) term "immune response" (GO:0006955).

| | |
|---|---|
| Hispid2B004960 | popy Class I histocompatibility antigen, A-1 alpha chain-like |
| Hispid2B005642 | interleukin-20 isoform X1 |
| Hispid2B007038 | histocompatibility 2, M region locus 1 |
| Hispid2B014408 | Ferritin, heavy polypeptide 1 |
| Hispid2B017133 | granzyme F |
| Hispid2B017337 | MHC class Ib antigen |
| Hispid2B018505 | carcinoembryonic antigen-related cell adhesion molecule 3-like isoform X2 |
| Hispid2B019905 | carcinoembryonic antigen-related cell adhesion molecule 3-like |
| Hispid2B028365 | cardiotrophin-1 isoform X1 |
| Hispid2B029895 | tetraspanin-4 isoform X1 |
| Hispid2B031114 | granzyme F |

Supplementary Table S3. **Immune response genes whose expression was not detected in cotton rat.**

Only 11 out of 224 selected genes represented in the GO term “immune response” (**Supp. Table S2**) were not detected in RNA pools prepared from multiple tissues of two cotton rat individuals. Their cotton rat identifiers are listed here, along with their assigned gene names.

| Geneid | naive_counts | hdm_counts | gene_name |
|---|---|---|---|
| Hispid2B031677 | 2,052 | 4,344 | CXCL9 |
| Hispid2B030611 | 5,856 | 28,686 | CXCR6_MOUSE |
| Hispid2B027351 | | | POK8_HUMAN |
| Hispid2B023916 | 3,014 | 2,842 | OASL2 |
| Hispid2B022981 | | | LORF2_MOUSE |
| Hispid2B021325 | 58 | 121 | LORF1_MOUSE |
| Hispid2B017969 | 657 | 547 | RSAD2 |
| Hispid2B015714 | 20,647 | 24,543 | GBP2 |
| Hispid2B014763 | 4,735 | 12,961 | GBP5 |
| Hispid2B014110 | 13,439 | 15,385 | IIGP1_MOUSE |
| Hispid2B013502 | | | VPK8_HUMAN |
| Hispid2B012902 | 195,934 | 228,405 | B2MG_SIGHI |
| Hispid2B011888 | 8,891 | 31,349 | GBP6_HUMAN |
| Hispid2B010441 | | | TAP1_RAT |
| Hispid2B010226 | 8,790 | 16,209 | IFIT1_MOUSE |
| Hispid2B007721 | 45,492 | 130,604 | CO7_PONAB |
| Hispid2B007382 | 7,433 | 10,962 | GBP4_MOUSE |
| Hispid2B001740 | 10,525 | 15,554 | KAL1L_HUMAN |
| Hispid2B001099 | 21,936 | 16,712 | MX2_MOUSE |

Supplementary Table S4. **Confirmation and annotation of differential gene expression upon RSV infection in cotton rats.** Sequence data of differentially expressed transcripts expressed in cotton rats that had been infected by RSV (Rajagopala et al. 2018) were downloaded. The cDNA sequences were aligned against the cotton rat genome assembly to support gene models. We aligned and counted RNA-seq reads from two cotton rats' tissue pools against these reference gene models as listed here.

GO:0002377 (immunoglobulin production)
GO:0003823 (antigen binding)
GO:0006955 (immune response)
GO:0006956 (complement activation)
GO:0071346 (cellular response to interferon-gamma.

Supplemental table S5A. **GO terms of immune genes duplicated in the cotton rat genome.**

GO:0002474 (antigen processing and presentation of peptide antigen via MHC class I)
GO:0003823 (antigen binding)
GO:0005132 (type I interferon receptor binding)
GO:0006952 (defense response)
GO:0006955 (immune response)
GO:0006958 (complement activation, classical pathway)
GO:0030881 (beta-2-microglobulin binding)
GO:0034987 (immunoglobulin receptor binding)
GO:0042742 (defense response to bacterium)
GO:0071352 (cellular response to interleukin-2)

Supplemental table S5B. **GO terms of immune genes duplicated in mouse genome.**

GO:0002504 (antigen processing and presentation of peptide or polysaccharide antigen via MHC class II)
GO:0006955 (immune response)
GO:0006958 (complement activation, classical pathway)
GO:0034987 (immunoglobulin receptor binding)
GO:0046635 (positive regulation of alpha-beta T cell activation)
GO:0060333 (interferon-gamma-mediated signaling pathway), GO:0003823 (antigen binding).

Supplemental table S5C. **GO terms of immune genes duplicated in human genome**.

| Repeat Class | mouse (mm10) no. elements | mouse (mm10) length (bp) | mouse (mm10) % genome sequence | cotton rat no. elements | cotton rat length (bp) | cotton rat % genome sequence |
|---|---|---|---|---|---|---|
| SINEs | 1,436,628 | 201,417,239 | 7.38 | 1,447,899 | 218,583,773 | 8.73 |
| LINEs | 690,722 | 535,566,664 | 19.61 | 545,998 | 387,470,083 | 15.47 |
| LTR elements | 789,257 | 323,579,563 | 11.85 | 664,809 | 245,117,281 | 9.79 |
| DNA elements | 151,750 | 30,012,106 | 1.10 | 78,418 | 13,741,388 | 0.55 |
| Unclassified | 20,696 | 9,886,221 | 0.36 | 216,668 | 56,989,845 | 2.28 |
| Small RNA | 19,494 | 1,624,349 | 0.06 | 5,251 | 1,181,467 | 0.05 |
| Satellites | 29,565 | 4,626,506 | 0.17 | 7,890 | 2,519,245 | 0.10 |
| Simple repeats | 1,351,337 | 70,761,142 | 2.59 | 853,766 | 40,264,455 | 1.61 |
| Low complexity | 157,021 | 10,005,493 | 0.37 | 103,761 | 5,819,139 | 0.23 |
| Total | 4,646,470 | 1,187,479,283 | 43.49 | 3,924,460 | 971,686,676 | 38.81 |

Supplementary Table S6. **Comparison of repetitive element counts in mouse vs. cotton rat genomes.**
